# The Renowned Flavor Compound Cinnamaldehyde Induces Sweet Taste by Targeting the Transmembrane Domain of T1R3 in the Sweet Taste Receptor

**DOI:** 10.1101/2025.04.03.643902

**Authors:** Tomoya Nakagita, Akihiro Itoigawa, Shinji Okada, Takumi Misaka

**Author notes:** Address correspondence to Takumi Misaka, Department of Applied Biological Chemistry, Graduate School of Agricultural and Life Sciences, The University of Tokyo, 1-1-1 Yayoi, Bunkyo-ku, Tokyo 113-8657, Japan.

## Abstract

Numerous flavor compounds are known to evoke sweet taste sensations, yet their direct interaction with the sweet taste receptor remains poorly understood. Traditionally, flavor-induced sweetness was attributed to aromatic properties rather than the taste qualities of these compounds. This study aims to elucidate whether the sweet taste receptor mediates flavor-induced sweet sensations by investigating the agonistic activities of 94 flavor compounds on cultured cells expressing the sweet taste receptor (T1R2/T1R3). The results demonstrated that cinnamaldehyde (CA) and *p*-methoxycinnamaldehyde (PMCA) activate the sweet taste receptor. These compounds not only induce the receptor response but also enhance receptor activity when combined with various sweeteners or sweet proteins. Due to PMCA’s structural similarity to lactisole, a well-known negative allosteric modulator (NAM) of the sweet taste receptor interacting with the transmembrane domain (TMD) of T1R3, CA and PMCA are hypothesized to interact with the T1R3 TMD as ago-PAMs (agonists and positive allosteric modulators). Mutational analyses confirmed that CA and PMCA interact with the T1R3 TMD, with distinct binding sites compared to lactisole. Additionally, because of structural parallels between PMCA and lactisole, we investigated the structure-activity relationships among 79 compounds to determine whether they function as ago-PAMs or NAMs. Most compounds acted as inhibitors, while those with specific planar structures acted as ago-PAMs. In conclusion, this study identified CA and PMCA as novel ago-PAMs that interact with the T1R3 TMD of the sweet taste receptor. These findings significantly advance our understanding of how flavor compounds influence sweet taste perception at the molecular level.

**Significance Statement:** Numerous flavor compounds evoke sweet taste sensations, yet their interaction with the sweet taste receptor is not well understood. This study identifies cinnamaldehyde (CA) and p-methoxycinnamaldehyde (PMCA) as activators of the sweet taste receptor, interacting with the transmembrane domain (TMD) of T1R3. CA and PMCA also act as positive allosteric modulators of other sweeteners (ago-PAMs). We discovered two binding sites in the T1R3 TMD, with CA and PMCA binding to different sites than lactisole, a known negative allosteric modulator. These findings identify CA and PMCA as novel ago-PAMs and significantly advance our understanding of how flavor compounds influence sweet taste perception at the molecular level.

## INTRODUCTION

Many flavor compounds are known to evoke a sweet taste and/or enhance sweetness when combined with various sweeteners (1–8). For example, cinnamaldehyde, a major compound in cinnamon tree essential oils, was evaluated in human sensory tests with nose clips and found to be appreciably sweeter than a 3% sucrose solution at concentrations of 0.5, 1.0, and 5.0 mg/mL (2). Additionally, the odors of strawberry, vanilla, almond, caramel, and pineapple have been reported to enhance the perceived sweetness of sucrose or aspartame solutions (5, 7, 9, 10). Constituents of these flavors, such as ethyl butyrate, furaneol, isoamyl acetate, and benzaldehyde, have also been reported to induce sweetness and/or have sweetness-enhancing effects (3, 11–14).

These flavor-induced sweet modulations can be potentially explained by at least two mechanisms: (I) the integration of olfactory and taste perception (i.e., cross-modal effects) and (II) direct taste modulation via taste receptors. Since the sweet-enhancing effects of many flavors can be abolished by preventing odorous compounds from reaching olfactory receptors by closing the external nares (1, 4, 7), most studies interpret these effects primarily as a result of cross-modal integration between odor and taste sensations within the brain. In other words, previous studies have conventionally attributed flavor-induced sweet enhancement to the sweet scent, not to the taste properties of these compounds.

However, since flavors in foods and beverages can directly act on taste receptors on the tongue, it is theoretically possible that orally ingested volatile flavors affect gustatory responses in the oral cavity. One purpose of our study was to investigate any agonistic activity of flavor compounds against the human sweet taste receptor. In mammals, sweet taste perception is mediated by a heterodimeric complex involving T1R2 and T1R3, both of which belong to class C G-protein-coupled receptors (GPCRs)(15, 16). T1Rs consist of a transmembrane domain (TMD) and a large extracellular domain (ECD). The ECD includes the cysteine-rich domain (CRD) and the Venus flytrap domain (VFTD) (17). The VFTD of T1R2 is known as the orthosteric binding site, which receives sugars and some artificial sweeteners (15, 18). The TMD of T1R3 is known as the allosteric binding site, which interacts with both positive and negative allosteric modulators (PAMs and NAMs). The sweeteners neohesperidin dihydrochalcone (NHDC) and cyclamate interact here and act as PAMs (19–21). Thus, agonists that interact with the TMD of class C GPCRs are generally called ago-PAMs. Lactisole and gymnemic acid are known to interact with the TMD to act as NAMs (22, 23). In our previous study, we investigated the detailed binding modes of lactisole and its derivative 2,4-DP and created docking models (24).

This study focused on volatile flavor compounds found in foods and beverages that have sweet tastes and/or sweet-enhancing effects, as validated empirically. These volatile flavor compounds possess a variety of chemical structures, including acids, alcohols, aldehydes, esters, ethers, and ketones. As noted, it is theoretically possible that flavor compounds could modulate the human taste receptor in the oral cavity, either directly or indirectly. To investigate the agonistic activity of flavor molecules, 94 chemicals were selected based on their sweet scent and/or reported taste properties (i.e., sweetness) from the journal “Perfumer & Flavorist.” Successful identification of a flavor or flavor mixture that can activate T1Rs below the odor threshold level would pave the way for novel applications of flavor molecules, such as sweet enhancers.

## RESULTS

### Identification of Flavor-Induced Activation of the Heterodimeric Sweet Taste Receptor, T1R2/T1R3

To elucidate whether flavors empirically described as sweet can activate the human sweet taste receptors, the agonistic activities of 94 flavor compounds were investigated using cultured cells expressing the sweet taste receptor(25). The agonistic activity was screened using a fluorescence plate reader assay to monitor the intracellular calcium concentration of the cultured cells stably expressing the wild-type (WT) sweet taste receptor (T1R2/T1R3) along with a chimeric G protein (Gα16gust44). These 94 flavor compounds, known for their sweet scent and/or taste, were classified into acids, alcohols, aldehydes, esters, ethers, and ketones (Supplemental Fig. 1). Among them, cinnamaldehyde (CA) and *p*-methoxycinnamaldehyde (PMCA) were identified as potential activators of the sweet taste receptor (Supplemental Fig. 1A). When tested at 1 mM, CA and PMCA increased calcium concentration in the sweet taste receptor-expressing cells to levels comparable to or greater than 1 mM aspartame, a known artificial sweetener (Supplemental Fig. 1A). The concentration dependency and specificity of their agonistic activity to the sweet taste receptor were confirmed (Fig 1A). Results showed that the flavor-induced effect on the sweet taste receptor was observed in a dose-dependent manner, for both of CA and of PMCA

**Figure. 1.**
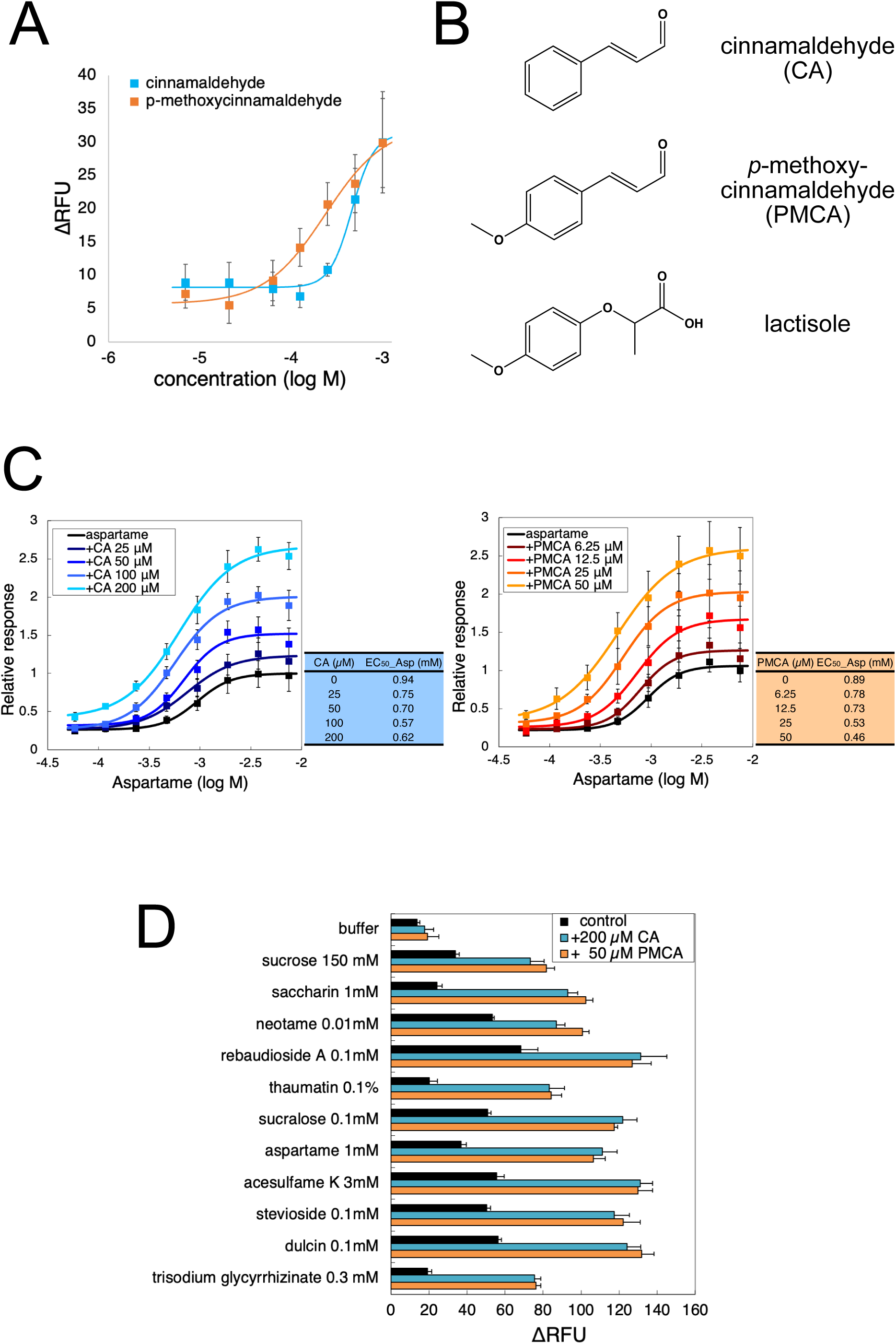
cinnamaldehyde and *p*-methoxycinnamaldehyde were obtained as agonist-positive allosteric modulator. After screening flavor compounds for their effects on the sweet taste receptor, cinnamaldehyde (CA) and *p*-methoxycinnamaldehyde (PMCA) were identified as agonist-positive allosteric modulators. (A) CA and PMCA elicit responses from the sweet taste receptor. The results of the flavor compound screening are shown in Supplemental Figure 1 and 2, and Supplemental Table 1. (B) Molecular structures of CA, PMCA, and the sweet taste inhibitor lactisole. (C) Dose-response curves of aspartame with varying concentrations of CA (left) and PMCA (right). Concentrations of CA added: 25 µM, 50 µM, 100 µM, and 200 µM; PMCA added: 6.25 µM, 12.5 µM, 25 µM, and 50 µM, respectively. EC_50_ values of aspartame under each condition are shown in the table. (D) Responses of the sweet taste receptor to sweeteners without and with 200 µM CA or 50 µM PMCA. All data correspond to mean ± SEM (n = 3-4).

### Cinnamaldehyde and *p*-Methoxycinnamaldehyde Potentiate the Sweet Taste Receptor Activities for Aspartame

Given the structural similarities between PMCA and lactisole (Fig. 1B), it was hypothesized that CA and PMCA interact with the T1R3 TMD similarly to lactisole(22, 24). To test this, the dose-response curves for aspartame were measured against the WT sweet taste receptor in the presence of various low concentrations of CA or PMCA (Fig. 1C). The administrated concentrations were 25 µM, 50 µM, 100 µM, and 200 µM for CA, and 6.25 µM, 12.5 µM, 25 µM and 50 µM PMCA respectively. Both CA and PMCA shifted the aspartame dose-response curve to the upper left in a dose-dependent manner. EC_50_ values for each dose-response curve are shown in Fig. 1C. The EC_50_ values of aspartame were reduced even with low concentrations of PMCA or CA, which showed only a minimal response on their own. Thus, these ligands are regarded as PAMs. To validate the specificity of CA- and PMCA-induced potentiation of T1R2/T1R3, receptor activity toward various sweeteners was measured using a cell-based assay in the presence of 200 µM CA or 50 µM PMCA, respectively (Fig. 1D). Both CA and PMCA significantly enhanced the potency of all tested sweeteners (i.e., sucrose, saccharin, neotame, rebaudioside A, thaumatin, sucralose, aspartame, acesulfame K, stevioside, dulcin, and trisodium glycyrrhizinate). The ΔRFU values for each sweetener increased from 1.6-fold to 4.1-fold with 0.2 mM CA, and from 1.9-fold to 4.2-fold with 0.05 mM PMCA, respectively. These results are similar to those seen with NHDC and cyclamate added to aspartame solution (21), indicating that CA and PMCA interact with T1R3 TMD.

### Identification of the Interaction Site of Cinnamaldehyde and *p*-Methoxycinnamaldehyde with the Sweet Taste Receptor

To identify the binding pocket of CA and PMCA, mutational analyses were conducted using 31 stable cell lines expressing point-mutated sweet taste receptors (Supplemental Table 2). Aspartame, which interacts with the VFTD of T1R2 (15), was used as a control ligand to ensure receptor function. EC_50_ values for aspartame are shown in Supplemental Table 2, with most mutants showing little EC_50_ fold change. However, S620A^2.52^, V721I^2.53^, and N737Q^5.47^ showed slight changes (colored in grey indicating a 5-fold change from EC_50__WT). The results for CA and PMCA are shown in Figure 2A, Supplemental Figure 3, and Supplemental Table 2. Mutants with significant fold changes in EC_50_ values for CA or PMCA are highlighted in Figure 2A, with dose-response curves shown in Figure 2B (mutants with no significant EC_50_ fold change are shown in Supplemental Figure 3). Mutants S620A^2.52^, F624L^2.56^, Q636A^3.32^, S729A^5.39^, and N737Q^5.47^ were significant for CA and PMCA interaction, especially Q636A^3.32^ and S729A^5.39^, where responses were significantly attenuated (colored in magenta for the crucial response reduction (Figure 2B)). PMCA-specific response reduction was observed in mutants S640A^3.36^, S726A^ECL2^, W727L^5.37^, and L782A^6.57^. In Q794N^7.32^, EC_50_ values for CA and PMCA shifted to more potent than WT (2x fold change colored in blue).

**Figure 2.**
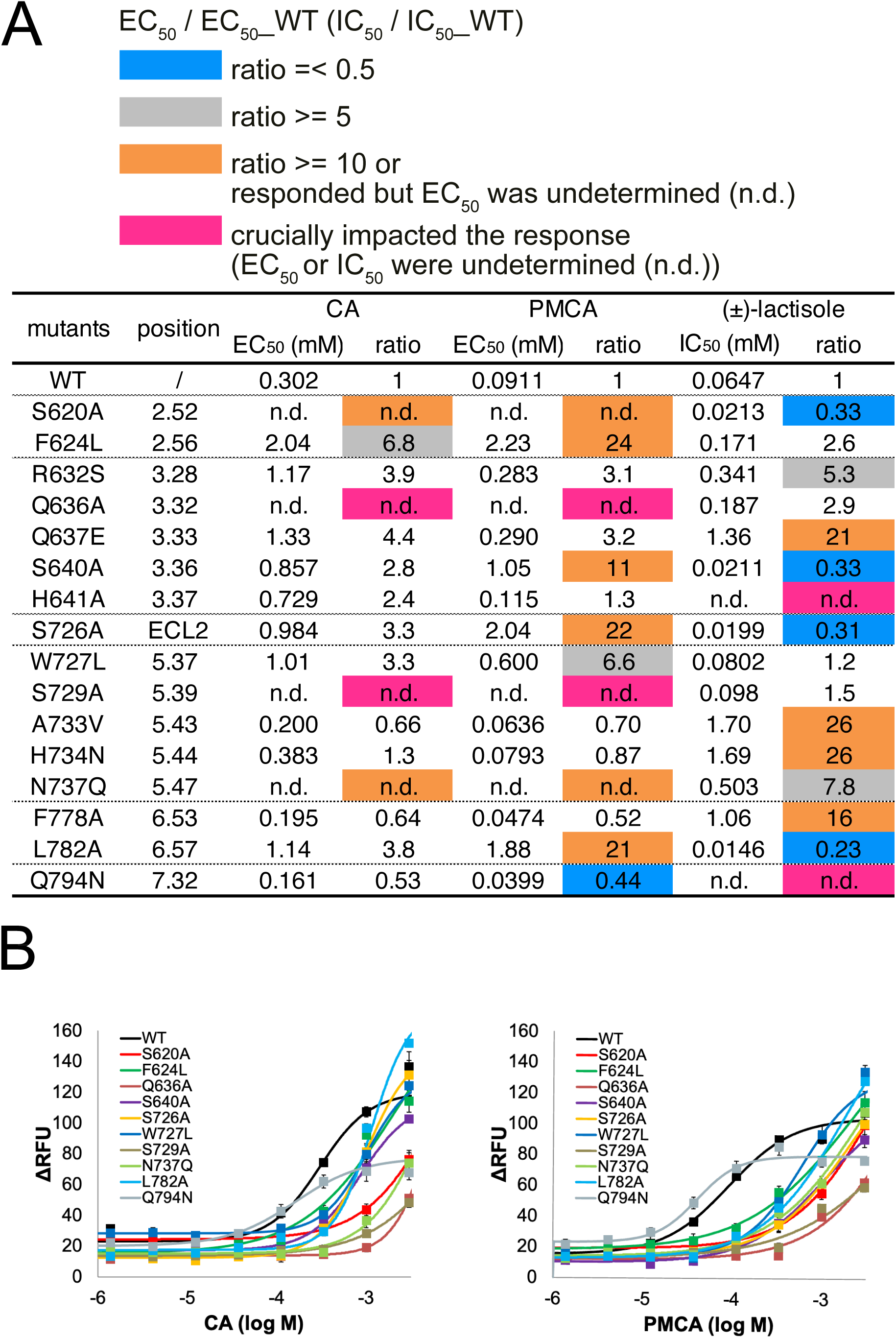
Results of mutational analyses of cinnamaldehyde, *p*-methoxycinnamaldehyde, and lactisole (A) Summary of mutation analyses. EC50 or IC50 values of each mutation are shown on the left side of each ligand column. The right-side column shows the ratio from the EC50 value of the wild type. Ratios over five are colored grey, over ten (including those where EC50 values couldn’t be determined but were expected) are colored orange. The most crucial mutants, for which EC50 or IC50 values couldn’t be determined, are shown in magenta. “n.d.” means not determined. Ratios of mutants less than 0.5 are shown in blue. All results analyzed in this study are also shown in Supplemental Tables 2 and 3. (B) Dose-response curves of mutants affecting the response to CA (left) and PMCA (right). Dose-response curves of all mutants for CA and PMCA are shown in Supplemental Figure 3. All data correspond to mean ± SEM (n = 3-4).

Previous studies showed that H641A^3.37^ and Q794N^7.32^ mutations reduced lactisole’s inhibitory activity (colored in magenta). Other critical mutations for lactisole were Q637E^3.33^, A733V^5.43^, H734N^5.44^, and F778A^6.53^ (colored in orange). Comparing mutants related to CA or PMCA and lactisole revealed that they were incompatible. In mutants S620A^2.52^, S640A^3.36^, S726A^ECL2^, L782A^6.57^, and Q794N^7.32^, when PAM or NAM lost its activity, the other gained it. These results strongly suggest two binding sites in T1R3 TMD: one for PAM and one for NAM.

In Supplemental Table 3, mutational analyses against NHDC and cyclamate showed Q636A^3.32^ and S729A^5.39^ were common mutants that reduced response against each ago-PAM. These residues are necessary to activate the sweet taste receptor.

### Creating a Docking Model of *p*-Methoxycinnamaldehyde with Active Form of T1R3 TMD Homology Model

Mutants S640A^3.36^, S726A^ECL2^, W727L^5.37^, and L782A^6.57^ showed a PMCA-specific response reduction, prompting docking simulations focused on these residues. The result of the PMCA docking model is shown in Figure 3A (also, docking result of CA is shown in Supplemental Figure 4). The left image shows the entire model of the T1R3 TMD. The upper right figure is a magnified view of the ligand-binding site, and the lower right figure is a 90° rotation of the upper right figure. Residues identified by mutational analyses are highlighted as same color. We docked the PMCA structure with its methoxy group near the W727^5.37^ and L782^6.57^ residues. most identified residues except S620^2.52^, S726^ECL2^ and N737^5.47^ are within 4 Å of PMCA.

**Figure 3.**
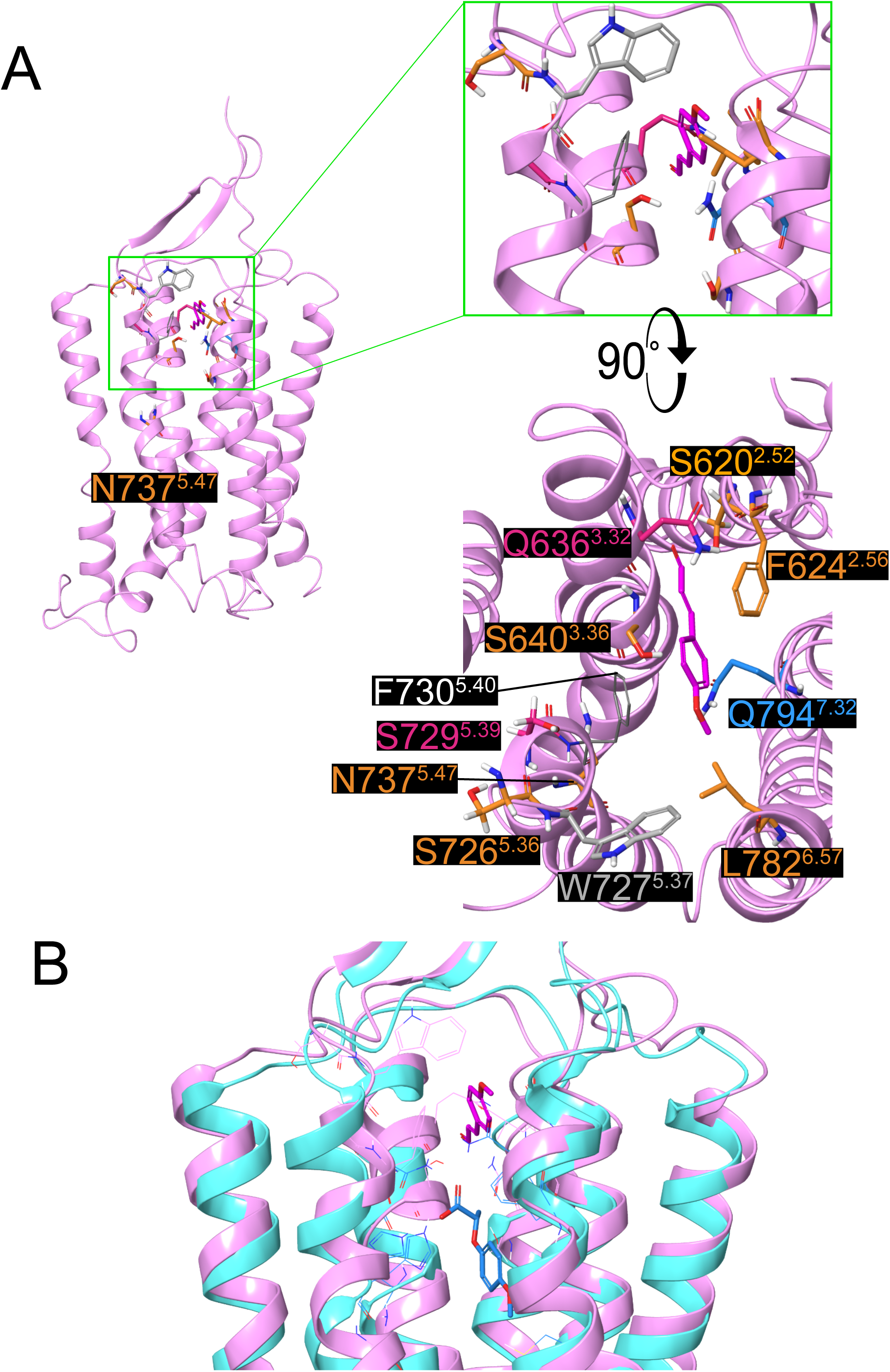
Results of docking simulation of *p*-methoxycinnamladehyde T1R3 TMD homology model based on the active form of the whole structure of mGluR5 (PDBID: 6N51). (A) Left: Overall view of T1R3-TMD and the binding site of PMCA. Upper right: Enlarged view of docked PMCA and surrounding residues. Lower right: 90° rotated view from upper right. Residue colors correspond to the mutational analyses in Figure 2. The docking result of CA is also shown in Supplemental Figure 4. (B) Superimposed view of PMCA docked active form T1R3 TMD (shown in pink ribbon) and lactisole docked apo-T1R3 TMD model (created by AlphaFold2, shown in cyan ribbon). Residues identified by mutational analyses are shown in their respective colors.

Docking simulations with other known PAMs, NHDC and cyclamate, were also conducted (Supplemental Figure 5A and 5B), with a superimposed view of PMCA, NHDC, and cyclamate shown in Supplemental Figure 5C. These PAMs share common binding sites, distinct from NAM binding sites. The lactisole docking model superimposed with the PMCA docking model is shown in Figure 3B. The active form of T1R3 is colored in pink ribbon, whereas the inactive form is in light blue ribbon. PMCA is colored magenta, and lactisole is colored blue.

### Investigating the Structure-Activity Relationships Between Lactisole and p-Methoxycinnamaldehyde

Given the similar molecular sizes of lactisole and PMCA, we investigated what defines a molecule as a PAM or NAM of the sweet taste receptor. Various concentrations of 79 compounds were added to 1 mM aspartame, and the threshold of activity as PAM or NAM was measured. The threshold was defined as the lowest concentration showing a p-value <= 0.05 in the student’s t-test. The list of 79 compounds is shown in Table 1, and the result summary is shown in Figure 4A. The horizontal axis shows the threshold concentration, and the vertical axis is ClogP (calculated log P), with higher values indicating more hydrophobicity. Grey bars indicate the ClogP area of the most potent PAMs. All measured data are shown in Supplemental Figure 5. Lactisole is compound 7, and PMCA is compound 79. Planar structures are shown as squares, partially planar structures as triangles, and flexible structures as circles. Compounds with a carboxyl group are colored blue, aldehydes red, alcohols green, ketones violet, and esters orange.

**Figure. 4.**
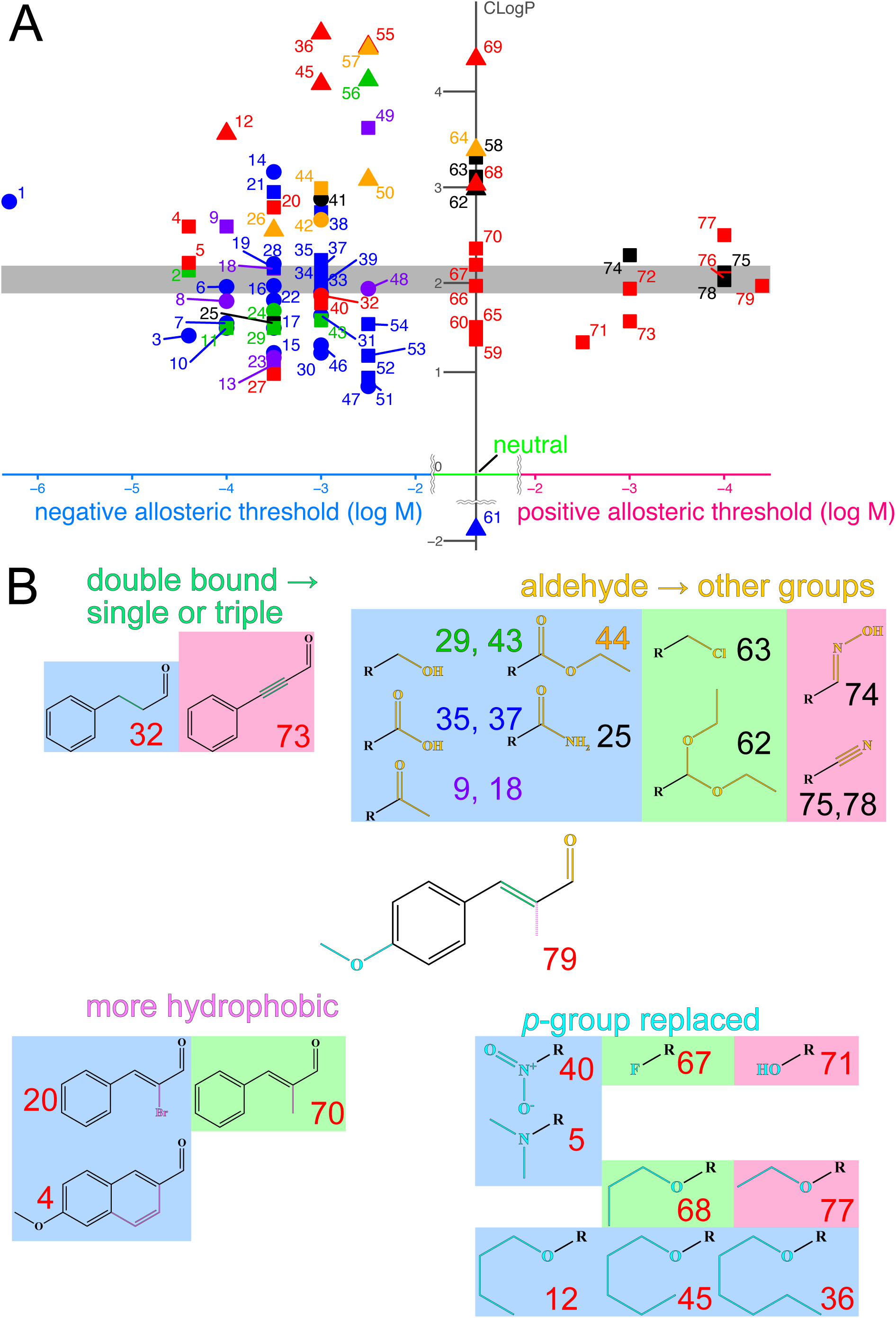
Investigation of structure-activity relationships between *p*-methoxycinnamaldehyde and lactisole All compounds were added to 1mM aspartame, and their PAM or NAM activities were measured. (A) Summary of the measurements for 79 compounds. The compound numbers correspond to Table 1. All raw data are shown in Supplemental Figure 6. The horizontal axis represents the threshold for raising PAM or NAM activities, while the vertical axis represents ClogP (calculated logP). The grey area indicates the ClogP range of the most effective PAMs. Compounds containing carboxyl groups are shown in blue, aldehyde groups in red, alcohol groups in green, ketone groups in violet, ester groups in orange, and others in black. Planar-shaped compounds are shown as squares, partially planar as triangles, and flexible shapes as circles. (B) Representative results of measured compounds. The colored parts of the PMCA molecular structure in the center indicate regions differing from the analogous structures measured in this study. Background colors correspond to NAM (blue), neutral (green), and PAM (red).

**Table 1.**
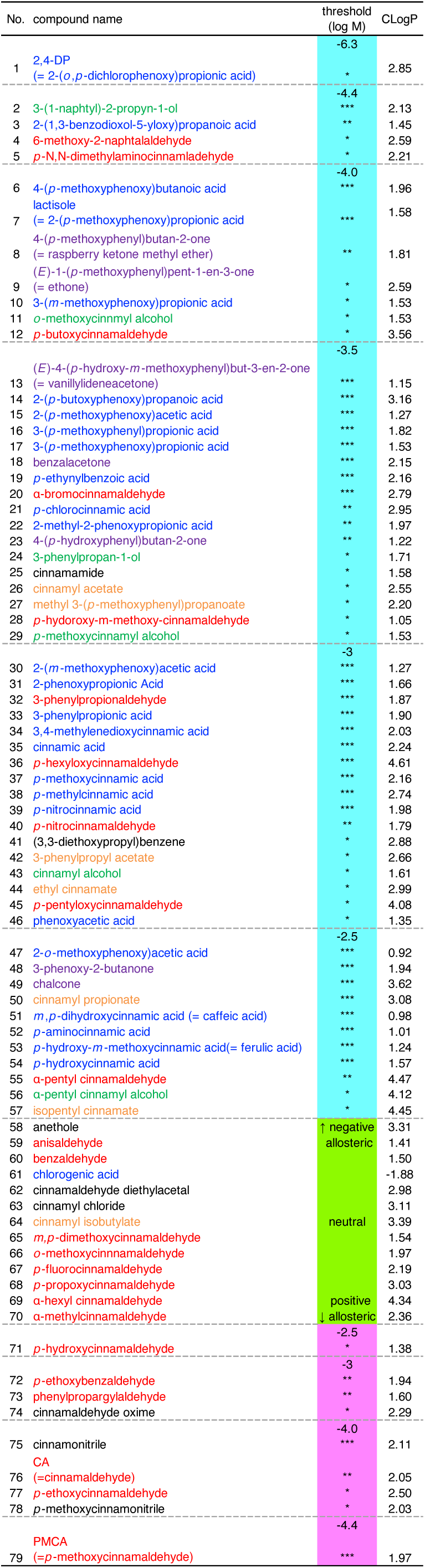
List of 79 compounds corresponding to Figure 4 colors of each compound also correspond to Figure 4. ClogP values calculated by ChemDraw 20.1 and the source of each compound is also shown here.

First, we examined the importance of the carbon chain between the aldehyde group and the aromatic ring (Fig. 4B, upper left). 3-phenylpropionaldehyde, compound No. **32**, which replaced the double bond of CA with a single bond, showed NAM activity at 1 mM, suggesting the importance of the planar structure of CA and PMCA. Phenylpropargyl aldehyde, compound **73**, acted as a PAM, supporting the importance of the planar structure for PAM activity.

Next, we investigated changing the aldehyde group of CA or PMCA to other groups. We examined cinnamic acid **35** and *p*-methoxycinnamic acid **37** as carboxyl groups, cinnamyl alcohol **43** and *p*-methoxycinnamyl alcohol **29** as alcohol groups, benzalacetone and 1-(*p*-methoxyphenyl)pent-1-en-3-one (ethone) **9** as ketone groups, cinnamonitrile **75** and *p*-methoxycinnamonitrile **78** as nitrile groups, cinnamamide **25** as an amide group, and ethyl cinnamate **44** as an ester group. Cinnamaldehyde diethylacetal **62**, cinnamyl chloride **63**, and cinnamaldehyde oxime **74** were also examined. Most groups, such as carboxyl, alcohol, ketone, ester, and amide, acted as NAMs, even if these compounds maintained the planar structure of cinnamaldehyde. However, only nitrile and oxime groups acted as PAMs. Cinnamaldehyde diethylacetal **62** and cinnamyl chloride **63** trended towards PAM activity (Supplemental Figure 5), but there was no significant difference from the response of 1 mM aspartame.

To further specify PAM activity, we measured compounds with a CA skeleton. α-methylcinnamaldehyde **70**, α-bromocinnamaldehyde **20** and 6-methoxy-2-naphthaldehyde **4** have more hydrophobic groups between the aldehyde and aromatic ring of CA. Compounds replacing the p-methoxy group of PMCA included *p*-hydroxycinnamaldehyde **71**, *p*-fluorocinnamaldehyde **67**, *p*-nitrocinnamaldehyde **40**, and *p*-N,N-dimethylaminocinnamaldehyde **5**. CA analogs with different lengths of alkoxyl groups were also examined: *p*-ethoxycinnamaldehyde **77**, *p*-propoxycinnamaldehyde **68**, *p*-butoxycinnamaldehyde **12**, *p*-pentyloxycinnamaldehyde **45**, and *p*-hexyloxycinnamaldehyde **36**.

Among these compounds, 6-methoxy-2-naphthaldehyde **4** and *p*-N,N-dimethylaminocinnamaldehyde **5** showed potent NAM activities, with thresholds of 37 µM, lower than lactisole (NAM threshold 110 µM). The ClogP values of these compounds were slightly higher than those of CA and PMCA. Other compounds generally appeared to have low affinity for T1R3 TMD if the ClogP was too low, and they were unable to enter the ligand pocket if it was too high. Although no more potent PAM than PMCA was identified, it was determined that PAMs interact with T1R3 TMD under very limited conditions: they must have a planar structure and aldehyde or nitrile groups, with small hydrophobic groups like p-methoxy groups.

## DISCUSSION

The primary question in this study was how flavor-induced sweet modulations could be explained. To find the answer, we examined whether 94 flavors actually elicit a response of the sweet taste receptor. The data generated from cell-based assays clarified that CA and PMCA had the potency to activate the sweet taste receptor at a dose of 1 mM (Supplemental Figures 1, 2 and Supplemental Table 1). The results support the hypothesis that, in some cases, sweetness or a sweet-enhancing effect is mediated by the sweet taste receptor, not by odor-taste integration in the brain. However, the remaining 92 flavors didn’t activate the sweet taste receptor, suggesting the existence of a cross-modal effect mechanism (i.e., an integration of odor and taste within the brain) since 1 mM was high enough for most flavor compounds to elicit an olfactory sensation on their own. Considering that in daily life, flavor-induced sweetness is sensed at doses less than 1 mM, this effect may be mediated by olfactory perception of “sweet scents” rather than gustatory input via the taste receptor.

CA and PMCA are detected not only as sweeteners (agonists) but also as ago-PAMs of the sweet taste receptor, interacting with T1R3 TMD. Indeed, the potentiating effect of both CA and PMCA was clearly detected with various sweeteners, even at low concentrations (e.g., 200 µM and 50 µM, respectively, in Fig. 1C), which are close to their olfactory threshold. Therefore, it is possible that when we enjoy cinnamon-flavored foods, they interact with the sweet taste receptor and provide sweetness. Additionally, using their properties as sweet taste enhancers, the development of sugar-reduced foods and beverages is a promising application.

In mutational analyses, mutants reducing CA’s and PMCA’s responses were different from those of NAM, lactisole. Docking simulations and these results strongly suggest that the ago-PAM and NAM are received by different positions of T1R3 TMD. There are few reports of different sites of action for PAMs and NAMs in this way, but they do exist. For example, in our previous study, methional, which interacts with T1R1 TMD of the umami taste receptor, is switchable between a PAM for the human and a NAM for the mouse umami taste receptor (26). In this case, the NAM binding mode of methional was expected to be in a deeper pocket of mouse T1R1 TMD. Furthermore, Robira *et al.* also mentioned that the binding sites of PAM and NAM of mGluR4 were different (27). In other cases, cryo-EM structures show that ADX55164 (ago-PAM of mGluR2 (PDBID 7MTR)) (28)and evocalcet (ago-PAM of CaSR (PDBID: 7DD7)) (29) occupy deeper pockets to shallow areas of the TMD binding site, as if they interact throughout the two binding sites we identified in this study (Supplemental Figure 7).

From supplemental Table 2 and supplemental Figure 4 and 5, CA and PMCA share important residues for interaction (especially Q636^3.32^ and S729^5.39^) and their binding site with NHDC and cyclamate. The shared area among these three ago-PAMs expands from TM3 to TM5 and TM6 under extracellular loop 2. However, in the model we created, CA and PMCA were placed a little farther from Ser726^ECL2^, Ser729^5.39^, and especially N737^5.47^. The N737Q^5.47^ mutation is too far from other candidate residues for CA and PMCA (Fig. 3A). This mutation might affect the receptor’s activation mechanism, such as conjugation with G-proteins, because the response of all sweeteners measured in this study, including aspartame as a control, decreased. Ser726^ECL2^ is at the end of extracellular loop 2, and Ser729^5.39^ is at the beginning of TM5. We hypothesized that these serine residues contribute to an increased OH-mediated hydrogen bonding network with NH in the main chain, which may stabilize a hydrophobic pocket comprising Trp727^5.37^, Phe730^5.40^ and Leu782^6.57^ in TMs 5 and 6. In addition, particularly for CA, residues extracted by mutational analyses contained almost no hydrophobic residues (with only a slight response reduction in Phe624^2.56^), despite CA’s structure being composed mostly of hydrophobic groups, except for the aldehyde group. Reviewing docking simulation results, the presence of Phe730^5.40^ also plays an important role in the interaction of these compounds with T1R3 TMD. In the mutational analyses of this study, we created F730L^5.40^ mutants according to previous studies (19, 20). In the F730L^5.40^ mutant, cyclamate and NHDC showed reduced activity against the sweet taste receptor. However, for CA or PMCA, it is possible that the L730^5.40^ mutation also engaged in the interaction.

Lastly, in five mutants (S620A^2.52^, S640A^3.36^, S726A^ECL2^, L782A^6.57^, and Q794N^7.32^), PMCA and lactisole showed interesting relationships: when PMCA activity reduced, lactisole activity increased, and vice versa. The structural similarity between these compounds may explain this phenomenon. Both PMCA and lactisole are not very potent as PAM or NAM, suggesting a probability that they might mistakenly enter each other’s pockets: PMCA into the NAM pocket and lactisole into the PAM pocket. These mutations likely make them more effective as PAMs and NAMs by reducing mistaken entry into the other pocket. These findings were revealed by comparing PMCA and lactisole, which have similar structures.

Finally, we investigated structure-activity relationships between PMCA and lactisole. Most compounds measured showed NAM activity or did not interact with the sweet taste receptor. Only a limited structure with a planar aldehyde or nitrile group of proper hydrophobicity showed PAM activity (Fig. 4A, B). In supplemental Table 3, we performed further mutational analyses against ethone **9** and *p*-N,N-dimethylaminocinnamaldehyde **5**. Both seem to interact with His641^3.37^, an essential residue for lactisole, but these interaction sites differ from that of lactisole and seem to have an intermediate mode interaction between PMCA and lactisole. The fact that H641^3.37^ is also reported to be the essential residue for gymnemic acid, another sweet-taste inhibitor known to have a different interaction mechanism form lactisole (23), suggesting that interaction on His641^3.37^ may be necessary to act as NAMs. Ethone **9** showed no crucial residues for interaction, suggesting it may interact with the PAM pocket while the ketone group interacts with His641^3.37^ instead of Q636^3.32^ due to the hydrophobicity of the ketone group. *p*-N,N-dimethylaminocinnamaldehyde **5** interacts with W727^5.37^, which can also interact with PMCA, but the aldehyde group interacts with His641^3.37^ instead of Q636^3.32^. In the case of *p*-N,N-dimethylaminocinnamaldehyde, the N,N-dimethyl group easily earns a positive charge. Because the aldehyde group is connected with it in resonance, it can turn into an enolate ion with a negative charge, causing interaction with His641^3.37^and exerting potent NAM activity.

Some commonly used flavor compounds, such as ethone **9** (sweet cherry and vanilla-like flavor) or raspberry ketone methyl ether **8** (sweet raspberry, rose, and cherry-like flavor), appear to have NAM activities in natural doses as flavors, despite having sweet scents. This inconsistency between taste and odor sensations is interesting.

In conlusion, we newly identified cinnamaldehyde and p-methocycinnamaldehyde as ago-PAMS of sweet taste receptor. We couldn’t identify a more potent ago-PAM than PMCA in this study. However, we gained many insights about binding mode of ago-PAMs; The common binding site among CA, PMCA, cyclamate and NHDC is different from that of NAM, lactisole. We hope that more potent ago-PAMs like ADX55164 in mGluR2 or evocalcet in CaSR, which can reduce the amount of sugar in food, are developed based on this information.

## MATERIALS AND METHODS

### Flavor compounds, sweet molecules and derivatives of lactisole and *p*-methoxycinnamaldehyde

The flavor compounds used in this study were purchased from venders (Tokyo Chemical Industry Co., Ltd., FUJIFILM Wako Pure Chemical Co., Kanto Chemical Co., Inc., and Sigma Aldrich). Sweeteners were purchased as follows: neohesperidin dihydrochalcone (NHDC), glycyrrhizic acid trisodium salt, and stevioside from Tokyo Chemical Industry Co., Ltd.; thaumatin, sucralose, aspartame, saccharin Na, and acesulfame K from FUJIFILM Wako Pure Chemical Co.; dulcin from Kanto Chemical Co., Inc.; neotame from NutraSweet Co. (Augusta, GA, USA); rebaudioside A from Morita Kagaku Kogyo (Osaka, Japan); sodium cyclamate from Sigma Aldrich; and sucrose from Nacalai Tesque (Kyoto, Japan). Lactisole and *p*-methoxycinnamaldehyde (PMCA) derivatives were purchased from Alfa Aeser (Thermo Fischer Scientific. Inc.), Across Organics (Thermo Fischer Scientific, Inc.), Kanto Chemical Co., Inc., NACALAI TESQUE, INC., Sigma-Aldrich Co. LLC, Tokyo Chemical Industry Co., Ltd. And FUJIFILM Wako Pure Chemical Co.. Some of the derivatives were synthesized. Calculation of each compound’s ClogP was performed by ChemDraw 20.1(Revvity, Inc, Waltham, MA, USA).

Ligands were once solubilized to DMSO in 1 M, and then diluted into the assay buffer at the desired concentrations. Ligands that used over 3 mM were directly solubilized by the assay buffer. The assay buffer was composed of 10 mM 4-(2-hydroxyethyl)-1-piperazineethanesulfonic acid (HEPES), 130 mM NaCl, 10 mM glucose, 5 mM KCl, 2 mM CaCl_2_, and 1.2 mM MgCl_2_ (pH adjusted to 7.4 using NaOH).

### Cell culture of the human sweet taste receptor expressing cells

As previously reported, Flp-In 293 cells that stably express T1R2/T1R3 together with Gα16gust44 were also used in this study (30). In addition of the stable cell line expressing the wild-type (WT) sweet taste receptor, 31 kinds of cell lines expressing its single point mutations in T1R3 TMD were also used (24). These cell lines were maintained in low-glucose (1.0 g/L) Dulbecco’s modified Eagle’s medium (Sigma Aldrich) with 10% fetal bovine serum (Thermo Fisher Scientific, Waltham, MA, USA).

### Measurement of cellular responses by fluorescence plate reader

Multiple data points and dose-response curves were generated from the data obtained in the cell-based assay using a FlexStation 3 (Molecular Devices). For the multiwell assays, cells were seeded in 96-well plates (CellBIND Black Plate, Corning Inc., Bedford, MA, USA) at approximately 70,000 cells per well. After 23h, cells were washed with the assay buffer before loading a calcium-indicator dye in the FLIPR Calcium 4 Assay Kit (Molecular Devices). Following that, the cells were incubated in two conditions in this study. In flavor assay (Figure 1), cells are incubated for 45 min at 27 _°_C to reduce background noises. In latter experiments (mutational assays and investigating structure-activity relationships of lactisole and PMCA), cells are incubated for 1h. at 37 _°_C to detect some mutants’ weak responses. Then, ligand solutions were administered into the wells automatically. Fluorescence changes (i.e., excitation at 485 nm and emission at 525 nm with a cutoff at 515 nm) were monitored at 2 s intervals. A 100 µL aliquot of the assay buffer supplemented with 2x ligands was added at 20 s and scanning was continued for an additional 100 s. The response of each well in the assay plate was determined by calculating ΔRFU (delta relative fluorescent units), as defined by the equation: (maximum fluorescent value) - (minimum fluorescent value). The data are shown as the mean ± SEM of the ΔRFU or relative response. EC_50_ values from dose-response data were calculated with Clampfit 10.4 (Axon Instruments, Foster City, CA, USA) using Hill’s equation.

#### Docking Simulations

For docking model of PMCA, cyclamate and NHDC: homology model of active formed sweet taste receptor was created by Prime program of Schrödinger suit 2019-4 (Schrödiner Inc., New York, NY, USA) based on the structure of whole structure of mGluR5 (PDB ID: 6N51) (31).

For docking model of (*S*)-lactisole: The model created by Alphafold2 was utilized (https://alphafold.ebi.ac.uk/entry/Q7RTX0). To create these models, we tried to utilize several templates, but these were the most satisfactory docking results. Some of residues’ rotamers were changed properly in each model.

The structure of NHDC was downloaded by pubchem, other compounds were drawn by ChemDraw 20.1. Each compound was energy-minimized and docked by MFmyPresto (FiatLux Corp., Tokyo, Japan). The docking poses were decided according to the results of mutational analyses we did.

## Abbreviations

ago-PAM: agonist-positive allosteric modulator
CA: cinnamaldehyde
CRD: cysteine-rich domain
ECD: extracellular domain
GPCR: G protein-coupled receptor
NAM: negative allosteric modulator
NHDC: neohesperidin dihydrochalcone
PAM: positive allosteric modulator
PMCA: *p*-methoxycinnamaldehyde
RFU: relative fluorescent units
TMD: transmembrane domain
VFTD: Venus flytrap domain
WT: wild-type.

## AUTHOR CONTRIBUTIONS

S.O. and M.T. contributed to the project planning and supervised the work. T.N. processed the experimental data, conducted analyses including *in silico* simulations, drafted the manuscript, and designed the figures. A.I. assisted with data analysis and figure design. All authors reviewed and approved the final manuscript.

## ACKOWLEDGEMENT

This work was partly supported by the Funding Program for Next Generation World-Leading Researchers from the Japan Society for the Promotion of Science (LS037), JSPS KAKENHI Grant number JP16H04918, and Takeda Science Foundation (all to T. M.).

JSPS KAKENHI Grant 23K05080 (to T. N.)

