## Supplemental Fig. 1 (+2-7), Supplemental Table 1 (+2,3) for "The Renowned Flavor Compound Cinnamaldehyde Induces Sweet Taste by Targeting the Transmembrane Domain of T1R3 in the Sweet Taste Receptor"

**Supplemental Figure 1. Screening of candidates as agonist for the sweet taste receptor from flavor compounds.**

(A)Calcium responses of Flp-In 293 cells stably expressing T1R2 and T1R3. ΔRFU values for each well are shown. 94 Flavors were applied to the receptor cells at a concentration of 1 mM. Arrows indicate obvious calcium responses with ΔRFU values greater than 20. (B) List of 94 flavor compounds used in this study. Well numbers indicate the location of each flavor on the 96-well assay plate in Supplemental Figure 1A.

**Supplemental Figure 2.** Chemical structures for the 94 flavor compounds used in Supplemental Figure 1.

Each compound is placed in the corresponding position in 96-well plate.

**Supplemental Table 1. List of flavor compounds and derivatives**

Summary of the profiles of the 94 flavor compounds, including names, chemical abstract registry numbers (CAS No.), FEMA numbers (if any), functional groups, and information about natural occurrence.

**Supplemental Figure 3.** Dose-response curves of mutants in T1R3 TMD not affecting $EC_{50}$ value fold change of CA or PMCA

(A) Dose-response curve of CA (B) Dose-response curve of CA PMCA. Each point represents mean ± SEM (n = 3-4).

Supplemental Table 2. Complete data of mutational analyses for wildtype and 31 mutants against aspartame and ago-PAMs.

Aspartame is measured as a positive control. Results for CA, PMCA, NHDC, and cyclamate are also shown. $EC_{50}$ values of each mutation are shown on the left side of each ligand column. The right-side column shows the ratio from the $EC_{50}$ value of the wild type. Ratios over five are colored grey, over ten (including those where $EC_{50}$ values couldn't be determined but were expected) are colored orange. The most crucial mutants, for which $EC_{50}$ values couldn't be determined, are shown in magenta. "n.d." means not determined. Ratios of mutants less than 0.5 are shown in blue.

Supplemental Table 3. Complete data of mutational analyses for wildtype and 31 mutants against NAMs.

Results for lactisole, ethone ((*E*)-1-(p-methoxyphenyl)pent-1-en-3-one) **9**, and *p*-N,N-dimethylaminoaldehyde **5** are shown. These numbers, shown in bold, correspond to Figure 4, Table 5, and Supplemental Figure 6. $IC_{50}$ values of each mutation are shown on the left side of each ligand column. The right-side column shows the ratio from the $IC_{50}$ value of the wild type. Ratios over five are colored grey, over ten are colored orange. The most crucial mutants, for which $IC_{50}$ values couldn't be determined, are shown in magenta. "n.d." means not determined. Ratios of mutants less than 0.5 are shown in blue.

Supplemental Figure 4. Comparison of docking results of cinnamaldehyde and *p*-metoxycinnamaldehyde

(A) Horizontal view of CA docked T1R3 TMD model. (B) Vertical view of CA docked T1R3 TMD model. Residues are shown as colors corresponding to Figure 2 and Supplemental Table2.

Supplemental Figure 5. Comparison of docking results of *p*-metoxycinnamaldehyde, cyclamate and NHDC.

(A) cyclamate, (B) NHDC, (C) superimposed with PMCA and these ago-PAMs. Horizontal views are shown on the left side, and vertical views on the right side. Residues are shown as colors corresponding to Supplemental Table2.

Supplemental Figure 6. Response data of all 79 compounds

All data correspond to mean ± SEM (n = 3-4). The compounds shown in Figure 4B have their numbers highlighted in red. P-values were calculated by Student's t-test. *: $p \leq 0.05$, **: $p \leq 0.01$, ***: $p \leq 0.005$.

Supplemental Figure 7. Superimposed view of ago-PAMs among class C GPCRs

PMCA of T1R3TMD is shown in magenta, ADX55164 of mGluR2 (from PDBID 7DD7) shown in yellow and evocalcet of CaSR (from 7MTR) is shown in blue.

supplemental Figure 1

A

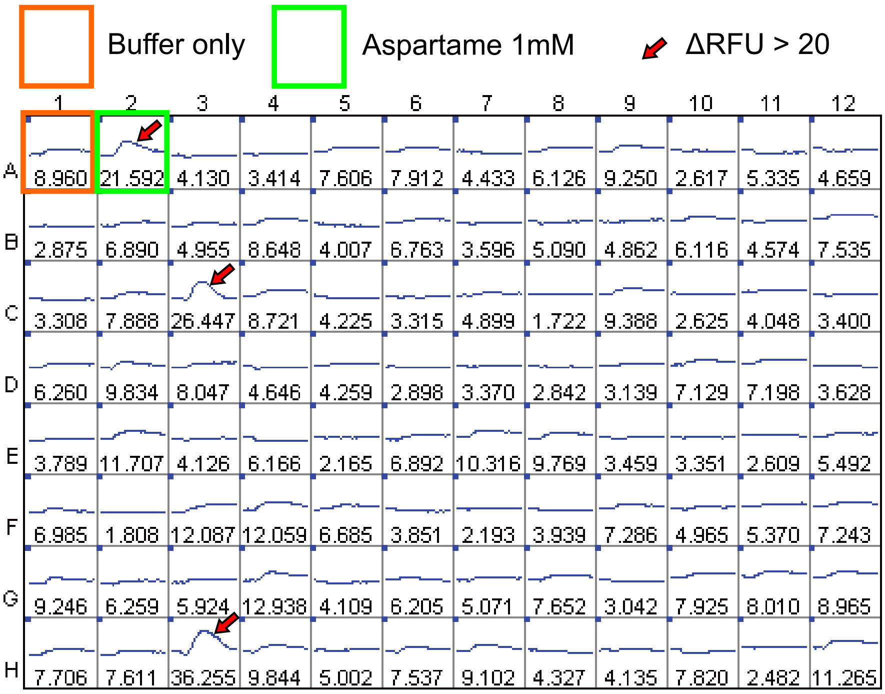


B

| well No. | fuctional group | compound name | well No. | fuctional group | compound name | well No. | fuctional group | compound name |
|---|---|---|---|---|---|---|---|---|
| A-1 | - | buffer | A-5 | ester | isoamylcinnamate | A-9 | ester | methyl-4-methyl valerate |
| B-1 | acid | vanillic acid | B-5 | ester | benzyl isobutyrate | B-9 | ester | γ-decalactone |
| C-1 | acid | tiglic acid | C-5 | ester | benzyl propionate | C-9 | ester | γ-heptalactone |
| D-1 | alcohol | cyclopentanol | D-5 | ester | allyl caproate | D-9 | ester | ethyl trans-3-hexenoate |
| E-1 | alcohol | guaiacol | E-5 | ester | ethyl anisate | E-9 | ester | methyl sorbate |
| F-1 | alcohol | eugenol | F-5 | ester | hexyl acetate | F-9 | ester | methyl cyclohexancarboxylate |
| G-1 | alcohol | isoeugenol | G-5 | ester | methyl anthranilate | G-9 | ester | anisyl acetate |
| H-1 | alcohol | trans-2-hexenol | H-5 | ester | δ-undecalactone | H-9 | ether | anethole |
| A-2 | - | aspartame | A-6 | ester | (-)-ethyl lactate | A-10 | ether | methyl chavicol |
| B-2 | alcohol | maltol | B-6 | ester | maltyl isobutyrate | B-10 | ether | limetol |
| C-2 | alcohol | vanillyl alcohol | C-6 | ester | ethyl phenylglycidate | C-10 | ether | vaniatrope |
| D-2 | alcohol | α-ionol | D-6 | ester | ethyl methylphenylglycidate | D-10 | ether | anisole |
| E-2 | alcohol | 2-phenylethyl alcohol | E-6 | ester | octalactone delta | E-10 | ether | methyleugenol |
| F-2 | alcohol | linalool | F-6 | ester | furfuryl pentanoate | F-10 | ether | methyl isoeugenol |
| G-2 | alcohol | ethyl maltol | G-6 | ester | phenylacetaldehyde | G-10 | ether | vanillin propylene glycol acetal |
| H-2 | alcohol | furfuryl alcohol | H-6 | ester | propyl caproate | H-10 | ether | vanillylidene acetone |
| A-3 | aldehyde | 4-ethoxybenzaldehyde | A-7 | ester | butyl phenylacetate | A-11 | ether | ethone |
| B-3 | aldehyde | benzaldehyde | B-7 | ester | ethyl trans-2-hexenoate | B-11 | ether | acetanisole |
| C-3 | aldehyde | cinnamaldehyde | C-7 | ester | methyl trans-3-hexenoate | C-11 | ether | allyl isovalerate |
| D-3 | aldehyde | heliotropine | D-7 | ester | δ-decalactone | D-11 | ketone | isophorone |
| E-3 | aldehyde | perillaldehyde | E-7 | ester | octyl butylate | E-11 | ketone | 2-furylpentyl ketone |
| F-3 | aldehyde | trans, trans-2,4-hexadienal | F-7 | ester | methyl trans-2-octenoate | F-11 | ketone | dihydrojasmone |
| G-3 | aldehyde | veratraldehyde | G-7 | ester | cyclohexyl propionate | G-11 | ketone | furaneol |
| H-3 | aldehyde | p-methoxycinnamaldehyde | H-7 | ester | allyl cyclohexyl propionate | H-11 | ketone | homosotolone |
| A-4 | aldehyde | p-methoxybenzaldehyde | A-8 | ester | cyclohexyl acetate | A-12 | ketone | p-methylacetophenone |
| B-4 | aldehyde | 5-methyl furfural | B-8 | ester | 6-methyl coumarin | B-12 | ketone | cyclotene |
| C-4 | aldehyde | 2-methyl-2-pentenal | C-8 | ester | menthalactone | C-12 | ketone | methyl heptadienone |
| D-4 | aldehyde | vanillin | D-8 | ester | ethyl benzoate | D-12 | ketone | α-damascone |
| E-4 | aldehyde | ethyl vanillin | E-8 | ester | γ-hexalactone | E-12 | ketone | homofuraneol |
| F-4 | aldehyde | α-amylcinnamaldehyde | F-8 | ester | anisyl formate | F-12 | ketone | acetophenone |
| G-4 | aldehyde | trans-2-heptenal | G-8 | ester | butyl butyrate | G-12 | ketone | 3-ethyl-2-hydroxy-2-cyclopente |
| H-4 | aldehyde | furfural | H-8 | ester | isobutyl phenylacetate | H-12 | ketone | acetoin |

supplemental Figure 2

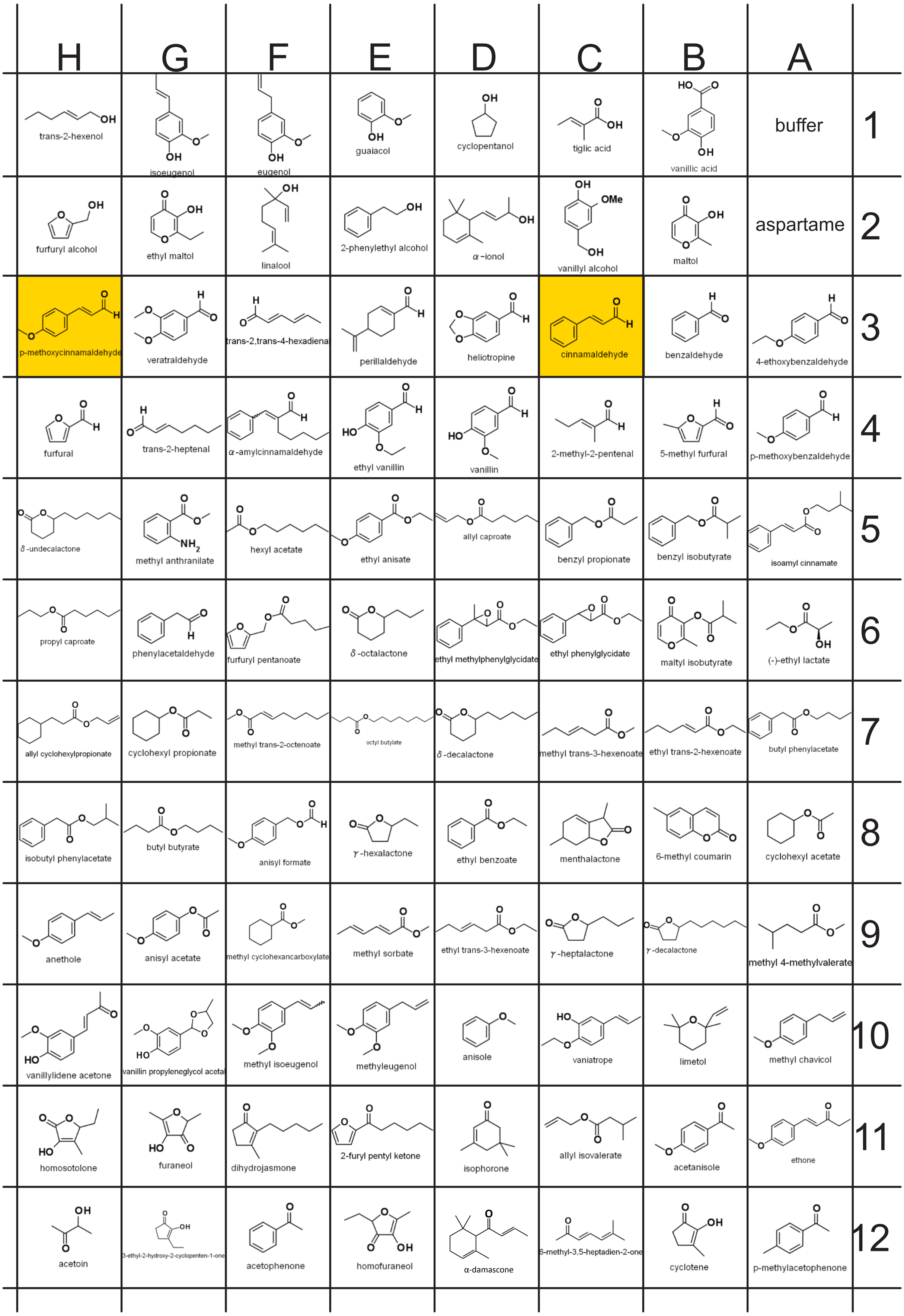

### supplemental Table 1

| position | name | CAS No. | FEMA No. | Functional group | | | Nature identical? | Natural occurrences |
|---|---|---|---|---|---|---|---|---|
| B-1 | vanillic acid | 121-34-6 | 3988 | acid | ether | aromatic | NI | |
| C-1 | tiglic acid | 80-59-1 | 3599 | acid | | | NI | celery leaves, |
| D-1 | cyclopentanol | 96-41-3 | | alcohol | | | NI | guava, peppermint oil, fish, beef |
| E-1 | guaiacol | 90-05-1 | 2532 | alcohol | ether | aromatic | NI | piper hancei, agarwood smoke, cistus, geranium, ginger flower |
| F-1 | eugenol | 97-53-0 | 2467 | alcohol | ether | aromatic | NI | apricot, banana, yuzu |
| G-1 | isoeugenol | 97-54-1 | 2468 | alcohol | ether | aromatic | NI | blueberry, guava fruit, tomato, clove bud |
| H-1 | trans-2-hexenol | 2305-21-7 | 2562 | alcohol | | | NI | green tea, kiwi fruit, tomato, strawberry, mango, apple |
| B-2 | maltol | 118-71-8 | 2656 | alcohol | ketone | | NI | coffee, strawberry, malt |
| C-2 | vanillyl alcohol | 498-00-0 | 3737 | alcohol | ether | | not found | |
| D-2 | α-ionol | 25312-34-9 | 3624 | alcohol | | | NI | osmanthus absolute |
| E-2 | 2-phenylethyl alcohol | 60-12-8 | 2858 | alcohol | aromatic | | NI | apple, apricot, banana, peach, pear, strawberry, cocoa, honey |
| F-2 | linalool | 8028-48-6 | 2635 | alcohol | | | NI | ho leaf, bitter orange flower, bois-de-rose, coriander |
| G-2 | ethyl maltol | 4940-11-8 | 3487 | alcohol | ketone | | not found | |
| H-2 | furfuryl alcohol | 98-000 | 2491 | alcohol | | | NI | cocoa, coffee, bread |
| A-3 | 4-ethoxybenzaldehyde | 10031-82-0 | 2413 | aldehyde | aromatic | | NI | black tea |
| B-3 | benzaldehyde | 100-52-7 | 2127 | aldehyde | aromatic | | NI | cherry pits, apricot pits |
| C-3 | cinnamaldehyde | 104-55-2 | 2286 | aldehyde | aromatic | | NI | cranberry, cassia oil |
| D-3 | heliotropine | 120-57-0 | 2911 | aldehyde | | | NI | vanilla, pepper, chicken |
| E-3 | perillaldehyde | 2111-75-3 | 3557 | aldehyde | | | NI | perilla frutescens, cumin, sweet orange |
| F-3 | trans, trans-2,4-hexadienal | 142-83-6 | 3429 | aldehyde | | | NI | kiwi fruit |
| G-3 | veratraldehyde | 120-14-9 | 3109 | aldehyde | ether | aromatic | NI | raspberry, peppermint oil |
| H-3 | p-methoxycinnamaldehyde | 24680-50-0 | 3567 | aldehyde | ether | aromatic | NI | cinnamon bark, cassia bark |
| A-4 | p-methoxybenzaldehyde | 123-11-5 | 2670 | aldehyde | ether | aromatic | NI | cranberry, fennel |
| B-4 | 5-methyl furfural | 620-02-0 | 2702 | aldehyde | | | NI | coffee, rum, bourbon whisky, cocoa, green tea, roasted almonds |
| C-4 | 2-methyl-2-pentenal | 623-36-9 | 3194 | aldehyde | | | NI | onion |
| D-4 | vanillin | 121-33-5 | 3107 | aldehyde | ether | aromatic | NI | vanilla, coffee, whisky, rum, nutmeg |
| E-4 | ethyl vanillin | 121-32-4 | 2464 | aldehyde | ether | aromatic | not found | |
| F-4 | α-amylcinnamaldehyde | 122-40-7 | 2061 | aldehyde | aromatic | | NI | black tea, soy bean |
| G-4 | trans-2-heptenal | 2463-63-0 | 3165 | aldehyde | | | NI | cranberry, asparagus, peanut roasted, potato chips, soy beans |
| H-4 | furfural | 98-01-1 | 2489 | aldehyde | | | NI | tamarind, averrhoa bilimbi fruit, mustard |
| A-5 | isoamylcinnamate | 7770-65-0 | 2063 | ester | | | NI | cinnamon |
| B-5 | benzyl isobutyrate | 103-28-6 | 2141 | ester | | | NI | mint, passion fruit, beer |
| C-5 | benzyl propionate | 122-63-4 | 2150 | ester | | | NI | plum |
| D-5 | allyl caproate | 123-68-2 | 2032 | ester | | | NI | pineapple, potato, mushroom |
| E-5 | ethyl anisate | 94-30-4 | 2420 | ester | ether | | NI | cassie blossom absolute |
| F-5 | hexyl acetate | 142-92-7 | 2565 | ester | | | NI | apple, banana, wine, mango |
| G-5 | methyl anthranilate | 134-20-3 | 2682 | ester | amine | | NI | strawberry, wine |
| H-5 | δ-undecalactone | 710-04-3 | 3294 | ester | lactone | | NI | butter |
| A-6 | (-)-ethyl lactate | 97-64-3 | 2440 | ester | | | NI | apple, apricot, pineapple |
| B-6 | maltyl isobutyrate | 65416-14-0 | 3462 | ester | | | not found | |
| C-6 | ethyl phenylglycidate | 121-39-1 | 2454 | ester | aromatic | | not found | |
| D-6 | ethyl methylphenylglycidate | 77-83-8 | 2444 | ester | aromatic | | not found | |
| E-6 | octalactone delta | 698-76-0 | 3214 | ester | lactone | | NI | rapsberry, butter, coconut, mango, wine |
| F-6 | furfuryl pentanoate | 36701-01-6 | 3397 | ester | | | NI | heated pork |
| G-6 | phenylacetaldehyde | 122-78-1 | 2874 | ester | | | NI | graipfruit juice, bilberry, tomato, white bread, cocoa |
| H-6 | propyl caproate | 626-77-7 | 2949 | ester | | | NI | apple juice |
| A-7 | butyl phenylacetate | 122-43-0 | 2209 | ester | | | NI | osmanthus concentrate |
| B-7 | ethyl trans-2-hexenoate | 27829-72-7 | 3675 | ester | | | NI | concord grape |
| C-7 | methyl trans-3-hexenoate | 2396-78-3 | 3364 | ester | | | NI | pineapple, mountain papaya |
| D-7 | δ-decalactone | 705-86-2 | 2361 | ester | lactone | | NI | peach, rapsberry, butter, rum, coconut |
| E-7 | octyl butylate | 110-39-4 | 2807 | ester | | | NI | strawberry |
| F-7 | methyl trans-2-octenoate | 2396-85-2 | 3712 | ester | | | NI | pear, pineapple |
| G-7 | cyclohexyl propionate | 6222-35-1 | 2354 | ester | | | not found | |
| H-7 | allyl cyclohexyl propionate | 2705-87-5 | 2026 | ester | | | not found | |
| A-8 | cyclohexyl acetate | 622-45-7 | 2349 | ester | | | NI | sauerkraut |
| B-8 | 6-methyl coumarin | 92-48-4 | 2699 | ester | lactone | aromatic | not found | |
| C-8 | menthalactone | 13341-72-5 | 3764 | ester | lactone | | NI | peppermint oil |
| D-8 | ethyl benzoate | 93-89-0 | 2422 | ester | | | NI | cranberry, black currant, peach, rapsberry, beer, rum, cocoa |
| E-8 | γ-hexalactone | 695-06-7 | 2556 | ester | lactone | | NI | milk |
| F-8 | anisyl formate | 122-91-8 | 2101 | ester | ether | aromatic | NI | vanilla beans |
| G-8 | butyl butyrate | 109-21-7 | 2186 | ester | | | NI | apple, orange, strawberry, mango |
| H-8 | isobutyl phenylacetate | 102-13-6 | 2210 | ester | | | NI | cocoa |
| A-9 | methyl-4-methyl valerate | 2412-80-8 | 2721 | ester | | | NI | pineapple |
| B-9 | γ-decalactone | 706-14-9 | 2360 | ester | | | NI | peach, butter, beer, rum, mango |
| C-9 | γ-heptalactone | 105-21-5 | 2539 | ester | lactone | | NI | green tea, asparagus, beer, strawberry, beef |
| D-9 | ethyl trans-3-hexenoate | 2396-83-0 | 3342 | ester | | | NI | melon, beer |
| E-9 | methyl sorbate | 689-89-4 | 3714 | ester | | | NI | pineapple, starfruit |
| F-9 | methyl cyclohexancarboxylate | 4630-82-4 | 3568 | ester | | | NI | vanilla |
| G-9 | anisyl acetate | 104-21-2 | 2098 | ester | ether | aromatic | NI | aristolochia asclepiadifolia root, ylang ylang |
| H-9 | anethole | 4180-23-8 | 2086 | ether | aromatic | | NI | cinnamon bark, nutmeg, dill, anise |
| A-10 | methyl chavicol | 140-67-0 | 2411 | ether | aromatic | | NI | anise oil |
| B-10 | limetol | 7392-19-0 | 3735 | ether | | | NI | geranium, lime, bitter orange leaf |
| C-10 | vaniatrope | 94-86-0 | 2922 | ether | aromatic | | not found | |
| D-10 | anisole | 100-66-3 | 2097 | ether | aromatic | | NI | truffle black, agarwood smoke, jasmine rose absolute |
| E-10 | methyleugenol | 93-15-2 | 2475 | ether | aromatic | | NI | pimento, bay, estragon, basil, laurel leaf |
| F-10 | methyl isoeugenol | 93-16-3 | 2476 | ether | aromatic | | NI | rose, basil, estragon, star anise |
| G-10 | vanillin propylene glycol acetal | 68527-74-2 | 3905 | ether | alcohol | | not found | |
| H-10 | vanillylidene acetone | 1080-12-2 | 3738 | ether | ketone | aromatic | not found | |
| A-11 | ethone | 104-27-8 | 2673 | ether | ketone | aromatic | not found | |
| B-11 | acetanisole | 100-06-1 | 2005 | ether | ketone | aromatic | NI | guava fruit, anise seed, celery seed |
| C-11 | allyl isovalerate | 2835-39-4 | 2045 | ether | | | not found | |
| D-11 | isophorone | 78-59-1 | 3553 | ketone | | | NI | grapefruit juice, cranberry, mushroom |
| E-11 | 2-furylpentyl ketone | 14360-50-0 | 3418 | ketone | | | not found | |
| F-11 | dihydrojasmone | 1128-08-1 | 3763 | ketone | | | not found | |
| G-11 | furaneol | 3658-77-3 | 3174 | ketone | lactone | | NI | coffee, grape, malt, pineapple, raspberry, strawberry |
| H-11 | homosotolone | 698-10-2 | 3153 | ketone | lactone | | NI | coffee |
| A-12 | p-methylacetophenone | 122-00-9 | 2677 | ketone | aromatic | | NI | mango, coocked cabbage, tomato |
| B-12 | cyclotene | 80-71-7 | 2700 | ketone | | | NI | coffee |
| C-12 | methyl heptadienone | 1604-28-0 | 3363 | ketone | | | NI | green tea |
| D-12 | α-damascone | 43052-87-5 | 3659 | ketone | | | NI | black tea |
| E-12 | homofuraneol | 27538-09-6 | 3623 | ketone | lactone | | NI | coffee, shoyu, strawberry, pineapple |
| F-12 | acetophenone | 98-86-2 | 2009 | ketone | aromatic | | NI | cocoa, beef, raspberry, peas |
| G-12 | 3-ethyl-2-hydroxy-cyclopentene-1-one | 21835-01-8 | 3152 | ketone | | | NI | coffee, tobacco |
| H-12 | acetoin | 513-86-0 | 2008 | ketone | alcohol | | NI | vinegar, wine, coffee, beer, cheddar |

supplemental Figure 3

A

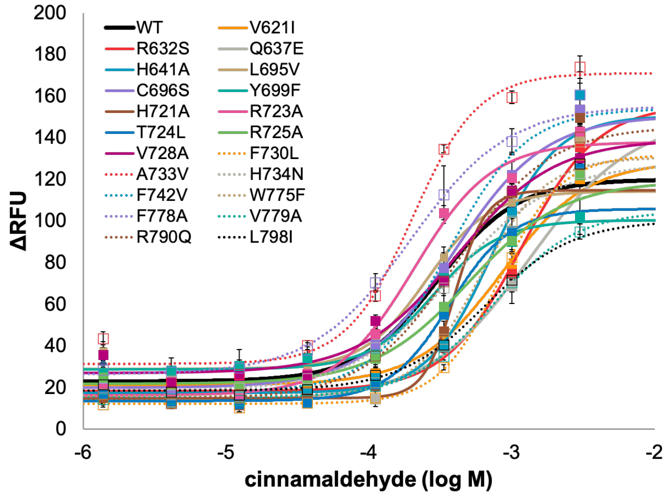


B

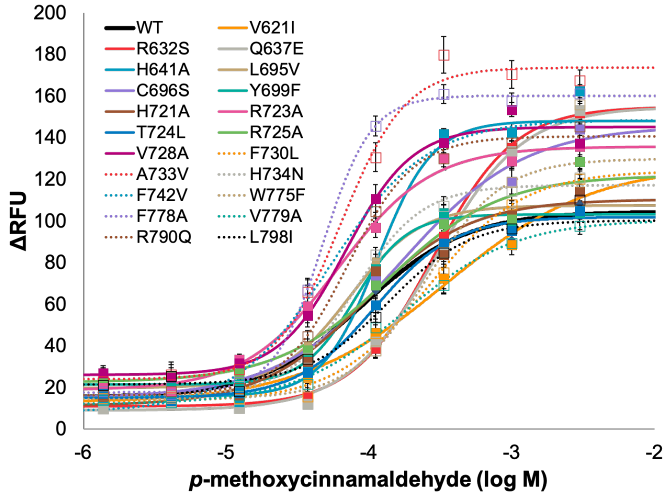

### supplemental Table 2

| mutants | position | aspartame | | CA **76** | | PMCA **79** | | cyclamate | | NHDC | |
|---|---|---|---|---|---|---|---|---|---|---|---|
| | | $EC_{50}$ (mM) | ratio | $EC_{50}$ (mM) | ratio | $EC_{50}$ (mM) | ratio | $EC_{50}$ (mM) | ratio | $EC_{50}$ (mM) | ratio |
| WT | / | 0.57 | 1 | 0.30 | 1 | 0.09 | 1 | 2.74 | 1 | 0.222 | 1 |
| S620A | 2.52 | 3.20 | 5.6 | n.d. | n.d. | n.d. | n.d. | 5.89 | 2.2 | 2.61 | 12 |
| V621I | 2.53 | 3.25 | 5.7 | 0.850 | 2.8 | 0.345 | 3.8 | 4.57 | 1.7 | 2.65 | 12 |
| F624L | 2.56 | 2.17 | 3.8 | 2.04 | 6.8 | 2.23 | 24 | n.d. | n.d. | 2.65 | 12 |
| R632S | 3.28 | 1.28 | 2.2 | 1.17 | 3.9 | 0.283 | 3.1 | 5.87 | 2.1 | 0.471 | 2.1 |
| Q636A | 3.32 | 1.16 | 2.0 | n.d. | n.d. | n.d. | n.d. | n.d. | n.d. | n.d. | n.d. |
| Q637E | 3.33 | 1.84 | 3.2 | 1.33 | 4.4 | 0.290 | 3.2 | n.d. | n.d. | 4.23 | 19 |
| S640A | 3.36 | 1.16 | 2.0 | 0.857 | 2.8 | 1.05 | 11 | 5.49 | 2.0 | 1.76 | 7.9 |
| H641A | 3.37 | 0.932 | 1.6 | 0.729 | 2.4 | 0.115 | 1.3 | n.d. | n.d. | 1.56 | 7.0 |
| L695V | 4.46 | 0.817 | 1.4 | 0.252 | 0.83 | 0.0732 | 0.80 | 2.14 | 0.78 | 0.178 | 0.80 |
| C696S | 4.47 | 0.746 | 1.3 | 0.393 | 1.3 | 0.168 | 1.8 | 8.65 | 3.2 | 0.434 | 2.0 |
| Y699F | 4.50 | 0.810 | 1.4 | 0.282 | 0.93 | 0.0793 | 0.87 | 5.67 | 2.1 | 1.62 | 7.3 |
| H721A | ECL2 | 1.85 | 3.2 | 0.399 | 1.3 | 0.0915 | 1.0 | 7.61 | 2.8 | 11.0 | 50 |
| R723A | ECL2 | 1.05 | 1.8 | 0.206 | 0.68 | 0.0606 | 0.67 | 7.47 | 2.7 | 0.128 | 0.58 |
| T724L | ECL2 | 1.38 | 2.4 | 0.367 | 1.2 | 0.110 | 1.2 | 7.30 | 2.7 | 0.367 | 1.7 |
| R725A | ECL2 | 1.98 | 3.5 | 0.497 | 1.6 | 0.137 | 1.5 | 4.92 | 1.8 | 0.229 | 1.0 |
| S726A | ECL2 | 2.09 | 3.6 | 0.984 | 3.3 | 2.04 | 22 | 15.6 | 5.7 | 0.738 | 3.3 |
| W727L | 5.37 | 1.89 | 3.3 | 1.01 | 3.3 | 0.600 | 6.6 | n.d. | n.d. | 2.27 | 10 |
| V728A | 5.38 | 0.985 | 1.7 | 0.408 | 1.4 | 0.0685 | 0.75 | 4.17 | 1.5 | 1.76 | 8.0 |
| S729A | 5.39 | 1.31 | 2.3 | n.d. | n.d. | n.d. | n.d. | n.d. | n.d. | n.d. | n.d. |
| F730L | 5.40 | 2.05 | 3.6 | 0.822 | 2.7 | 0.262 | 2.9 | n.d. | n.d. | n.d. | n.d. |
| A733V | 5.43 | 0.831 | 1.4 | 0.200 | 0.66 | 0.0636 | 0.70 | 3.28 | 1.2 | 0.130 | 0.6 |
| H734N | 5.44 | 0.935 | 1.6 | 0.383 | 1.3 | 0.0793 | 0.87 | n.d. | n.d. | 9.56 | 43 |
| N737Q | 5.47 | 3.05 | 5.3 | n.d. | n.d. | n.d. | n.d. | n.d. | n.d. | 9.20 | 42 |
| F742V | 5.52 | 0.510 | 0.89 | 0.362 | 1.2 | 0.0533 | 0.59 | 2.07 | 0.76 | 0.233 | 1.1 |
| W775F | 6.50 | 1.60 | 2.8 | 0.602 | 2.0 | 0.254 | 2.8 | 6.70 | 2.4 | 0.613 | 2.8 |
| F778A | 6.53 | 0.587 | 1.0 | 0.195 | 0.64 | 0.0474 | 0.52 | 11.4 | 4.2 | 1.45 | 6.5 |
| V779A | 6.54 | 1.23 | 2.1 | 0.743 | 2.5 | 0.176 | 1.9 | 2.94 | 1.1 | 0.240 | 1.1 |
| L782A | 6.57 | 1.21 | 2.1 | 1.14 | 3.8 | 1.88 | 21 | n.d. | n.d. | 1.25 | 5.6 |
| R790Q | 7.28 | 0.628 | 1.1 | 0.410 | 1.4 | 0.0705 | 0.77 | n.d. | n.d. | 0.134 | 0.60 |
| Q794N | 7.32 | 0.570 | 1.0 | 0.161 | 0.53 | 0.0399 | 0.44 | 3.13 | 1.1 | 0.571 | 2.6 |
| L798I | 7.36 | 1.07 | 1.9 | 0.653 | 2.2 | 0.136 | 1.5 | 4.04 | 1.5 | 0.296 | 1.3 |

### supplemental Table 3

| mutants | position | (±)-lactisole **7** | | ethone **9** | | *p*-N,N-dimethylamino-cinnamaldehyde **5** | |
|---|---|---|---|---|---|---|---|
| | | $IC_{50}$ (mM) | ratio | $IC_{50}$ (mM) | ratio | $IC_{50}$ (mM) | ratio |
| WT | / | 0.0647 | 1 | 0.118 | 1 | 0.09 | 1 |
| S620A | 2.52 | 0.0213 | 0.33 | 0.0565 | 0.48 | 0.06 | 0.73 |
| V621I | 2.53 | 0.0449 | 0.70 | 0.0556 | 0.47 | 0.06 | 0.73 |
| F624L | 2.56 | 0.171 | 2.6 | 0.0467 | 0.39 | 0.10 | 1.1 |
| R632S | 3.28 | 0.341 | 5.3 | 0.137 | 1.2 | 0.12 | 1.4 |
| Q636A | 3.32 | 0.187 | 2.9 | 0.0228 | 0.19 | 0.10 | 1.2 |
| Q637E | 3.33 | 1.36 | 21 | 0.329 | 2.8 | 0.21 | 2.5 |
| S640A | 3.36 | 0.0211 | 0.33 | 0.0398 | 0.34 | 0.09 | 1.0 |
| H641A | 3.37 | n.d. | n.d. | 0.626 | 5.3 | 2.12 | 24 |
| L695V | 4.46 | 0.0485 | 0.75 | 0.130 | 1.1 | 0.10 | 1.1 |
| C696S | 4.47 | 0.106 | 1.6 | 0.121 | 1.0 | 0.09 | 1.1 |
| Y699F | 4.50 | 0.294 | 4.6 | 0.113 | 0.96 | 0.34 | 3.9 |
| H721A | ECL2 | 0.0745 | 1.2 | 0.103 | 0.87 | 0.10 | 1.2 |
| R723A | ECL2 | 0.164 | 2.5 | 0.369 | 3.1 | 0.14 | 1.6 |
| T724L | ECL2 | 0.097 | 1.5 | 0.140 | 1.2 | 0.09 | 0.98 |
| R725A | ECL2 | 0.0601 | 0.93 | 0.0658 | 0.56 | 0.07 | 0.83 |
| S726A | ECL2 | 0.0199 | 0.31 | 0.099 | 0.83 | 0.07 | 0.80 |
| W727L | 5.37 | 0.0802 | 1.2 | 0.126 | 1.1 | n.d. | n.d. |
| V728A | 5.38 | 0.292 | 4.5 | 0.106 | 0.90 | 0.10 | 1.2 |
| S729A | 5.39 | 0.098 | 1.5 | 0.152 | 1.3 | 0.15 | 1.7 |
| F730L | 5.40 | 0.0775 | 1.2 | 0.0533 | 0.45 | 0.08 | 0.87 |
| A733V | 5.43 | 1.70 | 26.3 | 0.680 | 5.7 | 0.58 | 6.7 |
| H734N | 5.44 | 1.69 | 26.1 | 0.354 | 3.0 | 0.25 | 2.9 |
| N737Q | 5.47 | 0.503 | 7.8 | 0.0513 | 0.43 | 0.05 | 0.63 |
| F742V | 5.52 | 0.126 | 1.9 | 0.362 | 3.1 | 0.14 | 1.6 |
| W775F | 6.50 | 0.0741 | 1.1 | 0.0860 | 0.73 | 0.12 | 1.3 |
| F778A | 6.53 | 1.06 | 16.4 | 0.473 | 4.0 | 0.28 | 3.2 |
| V779A | 6.54 | 0.158 | 2.4 | 0.142 | 1.2 | 0.13 | 1.5 |
| L782A | 6.57 | 0.0146 | 0.23 | 0.0545 | 0.46 | 0.04 | 0.51 |
| R790Q | 7.28 | 0.200 | 3.1 | 0.280 | 2.4 | 0.16 | 1.8 |
| Q794N | 7.32 | n.d. | n.d. | 0.278 | 2.3 | 0.08 | 0.92 |
| L798I | 7.36 | 0.144 | 2.2 | 0.0653 | 0.55 | 0.05 | 0.53 |

supplemental Figure 4

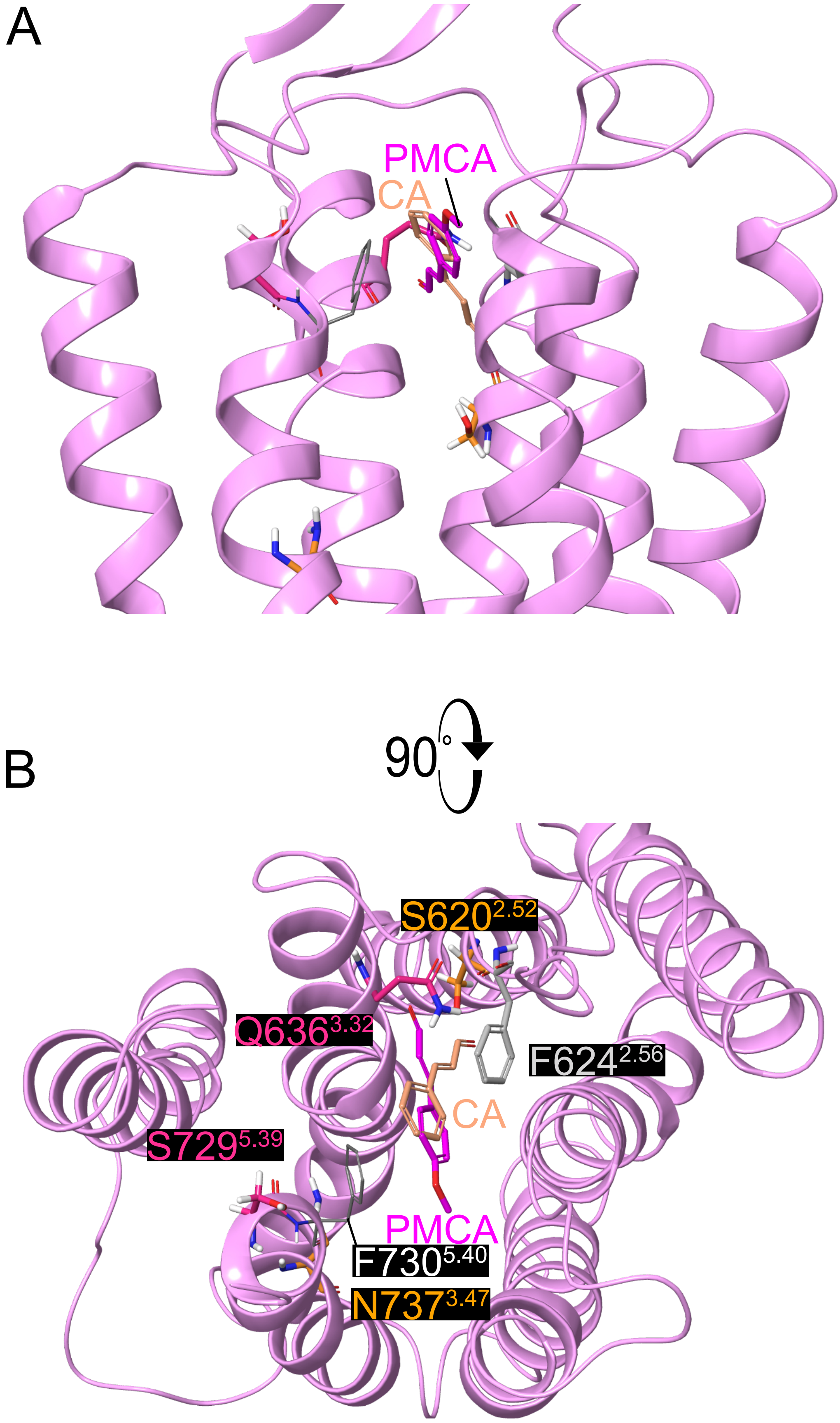

supplemental Figure 5

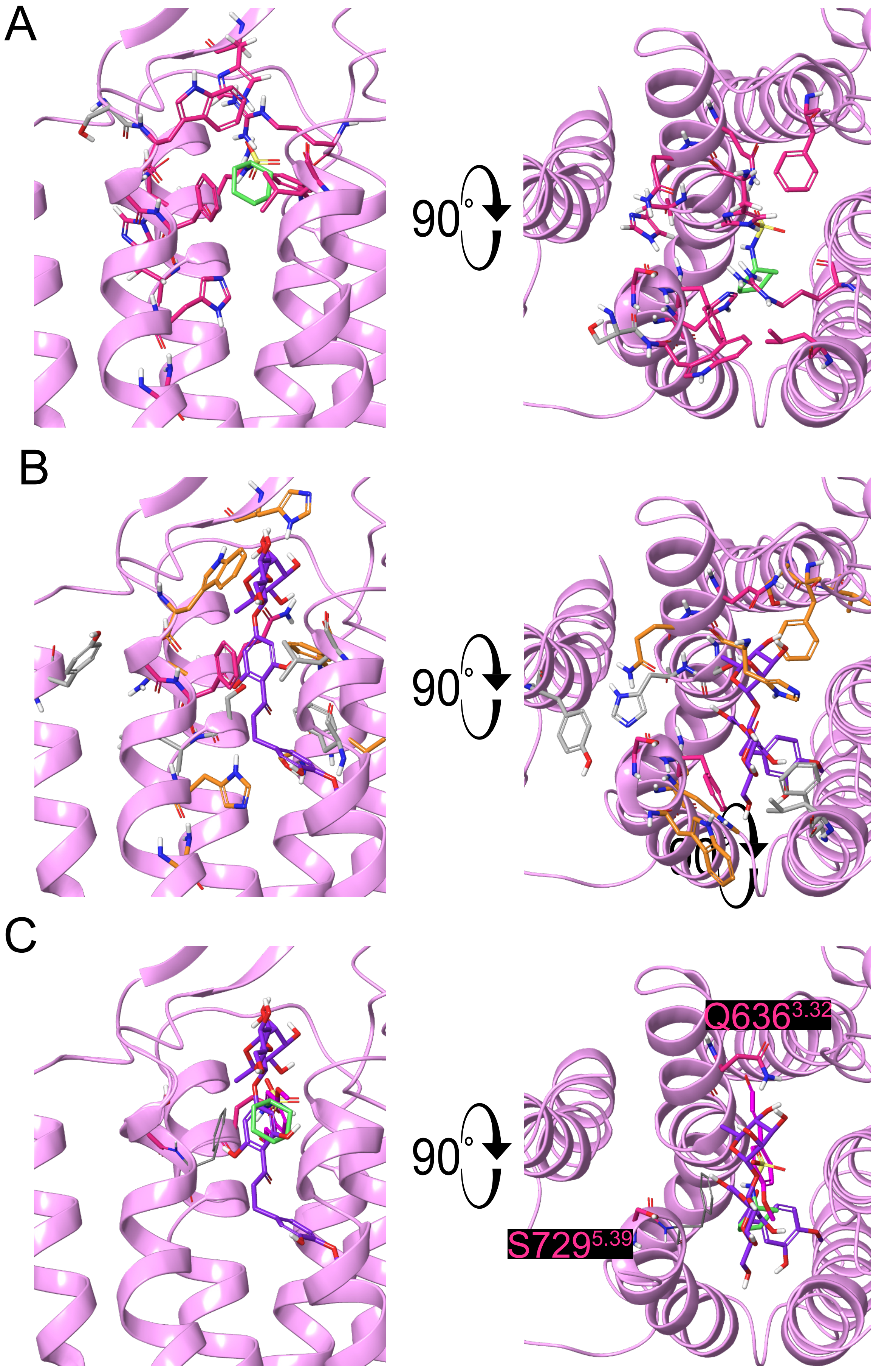

### supplemental Figure 6

#### Negative allosteric threshold -6.3

**1** **2-(*o*,*p*-dichlorophenoxy)propionic acid (2,4-DP)**

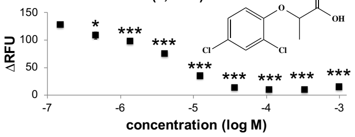


#### Negative allosteric threshold -4.4

**2** **3-(1-naphtyl)-2-propyn-1-ol**

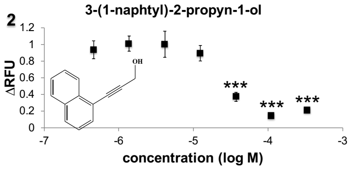


**3** **2-(1,3-benzodioxol-5-yloxy)propionic acid**

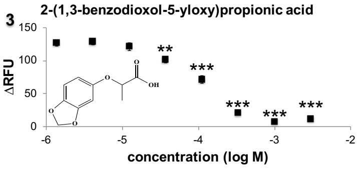


**4** **6-methoxy-2-naphtalaldehyde**

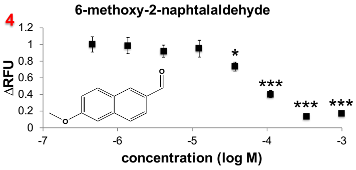


**5** ***p*-N,N-dimethylaminocinnamaldehyde**

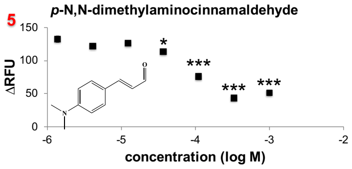


#### Negative allosteric threshold -4.0

**6** **4-(*p*-methoxyphenoxy)butanoic acid**

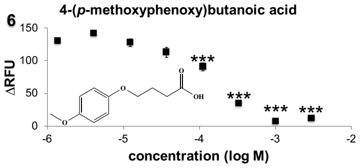


**7** **2-(*p*-methoxyphenoxy)propionic acid (lactisole)**

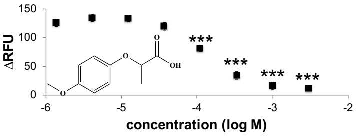


**8** **4-(*p*-methoxyphenylbutan)-2-one (raspberry ketone methyl ether)**

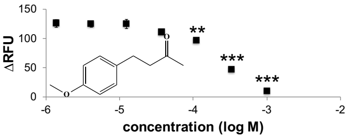


**9** **(*E*)-1-(*p*-methoxyphenyl)pent-1-en-3-one (ethone)**

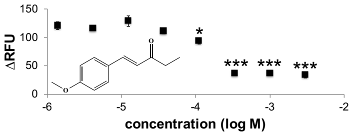


**10** **3-(*m*-methoxyphenoxy)propionic acid**

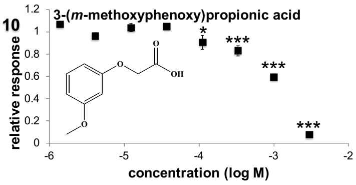


**11** ***o*-methoxycinnamyl alcohol**

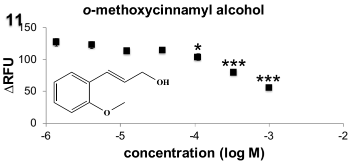


**12** ***p*-butoxycinnamaldehyde**

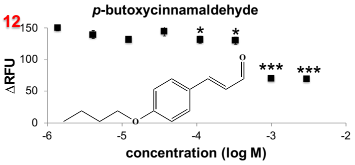


#### Negative allosteric threshold -3.5

**13** **(*E*)-4-(*p*-hydroxy-*m*-methoxyphenyl)but-3-en-2-one (vanillilidene acetone)**

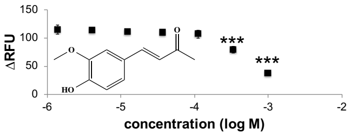


**14** **2-(*p*-butoxyphenoxy)propanoic acid**

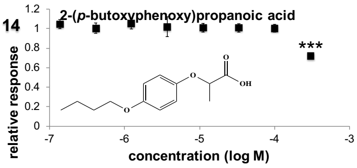


**15** **2-(*p*-methoxyphenoxy)acetic acid**

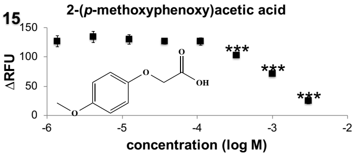


**16** **3-(*p*-methoxyphenyl)propionic acid**

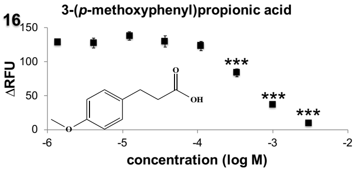


**17** **3-(*p*-methoxyphenoxy)propionic acid**

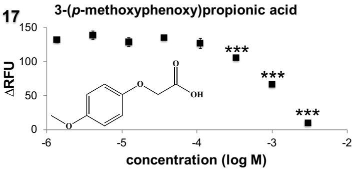


**18** **benzalacetone**

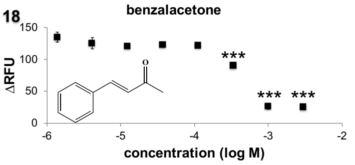


**19** **4-ethynylbenzoic acid**

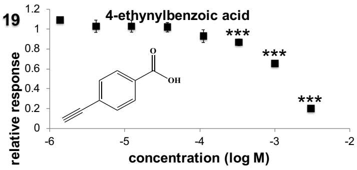


**20** **α-bromocinnamaldehyde**

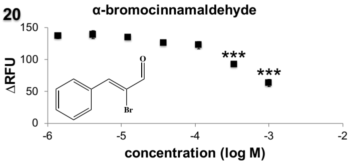


**21** ***p*-chlorocinnamic acid**

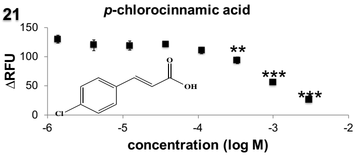

### supplemental Figure 6

#### Negative allosteric threshold -3.5 (continued)

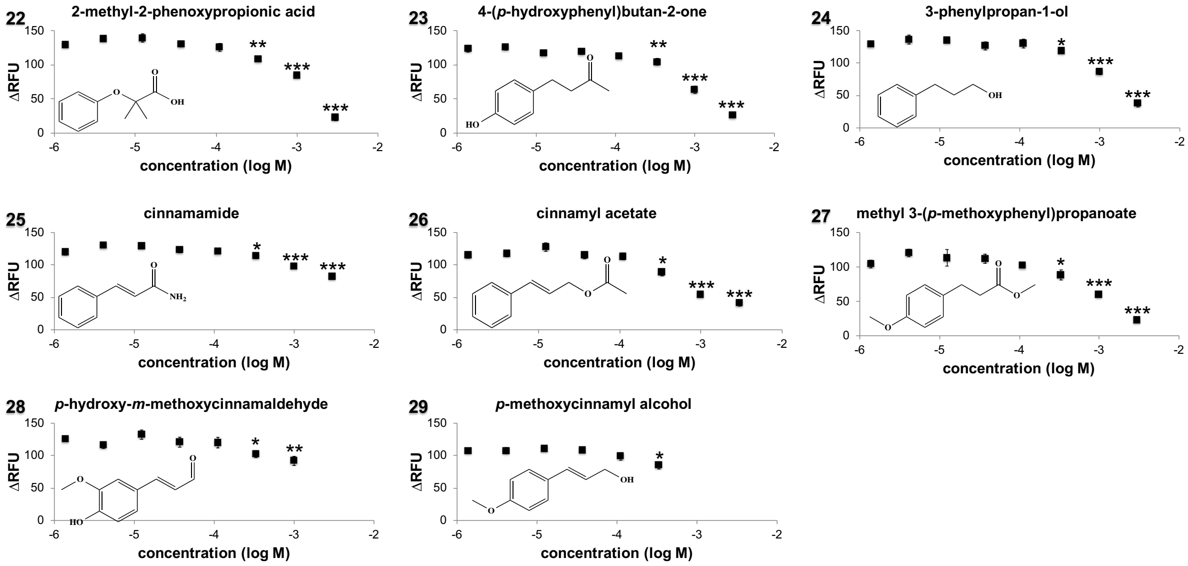


#### Negative allosteric threshold -3.0

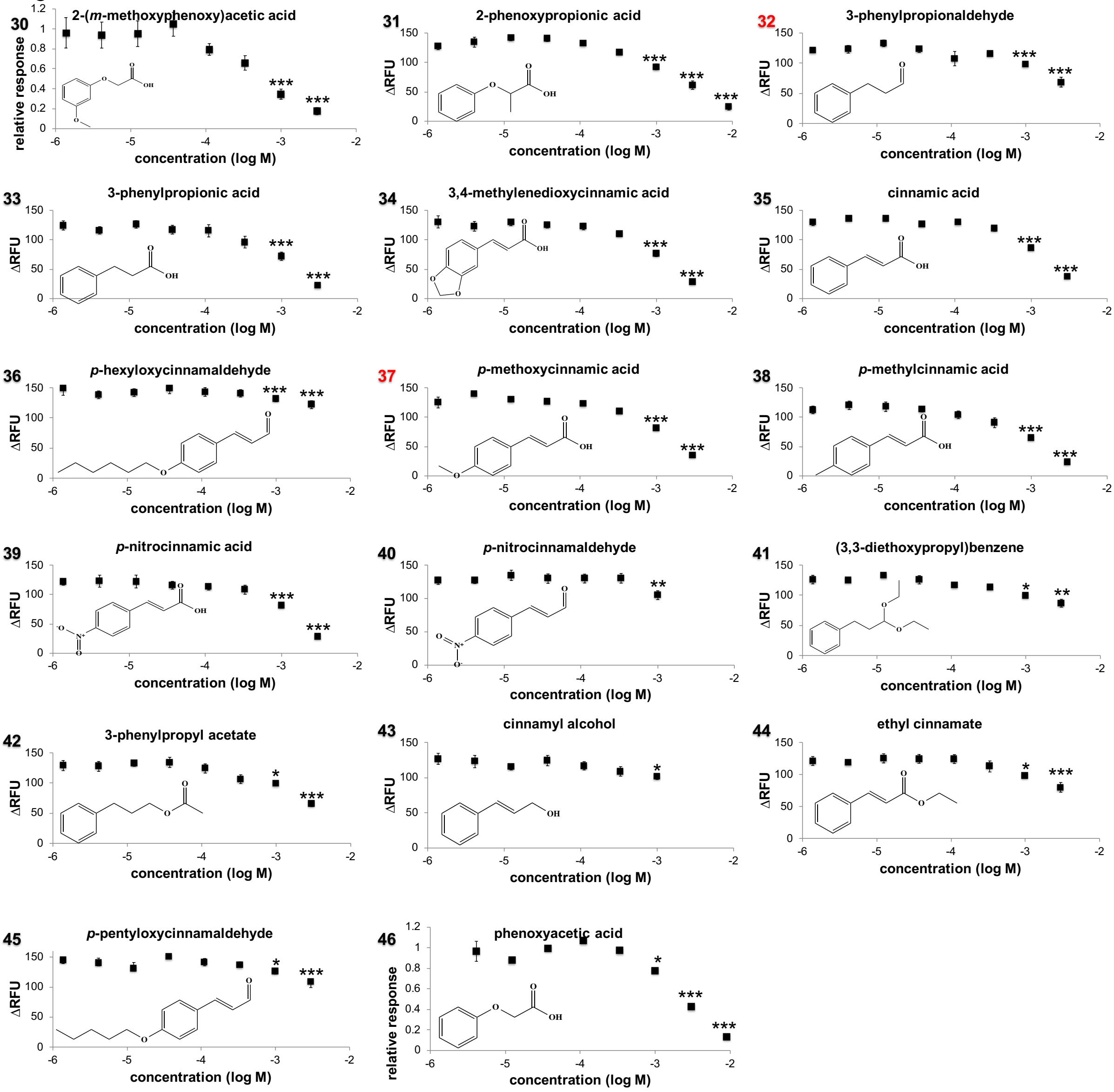

### supplemental Figure 6

#### Negative allosteric threshold -2.5

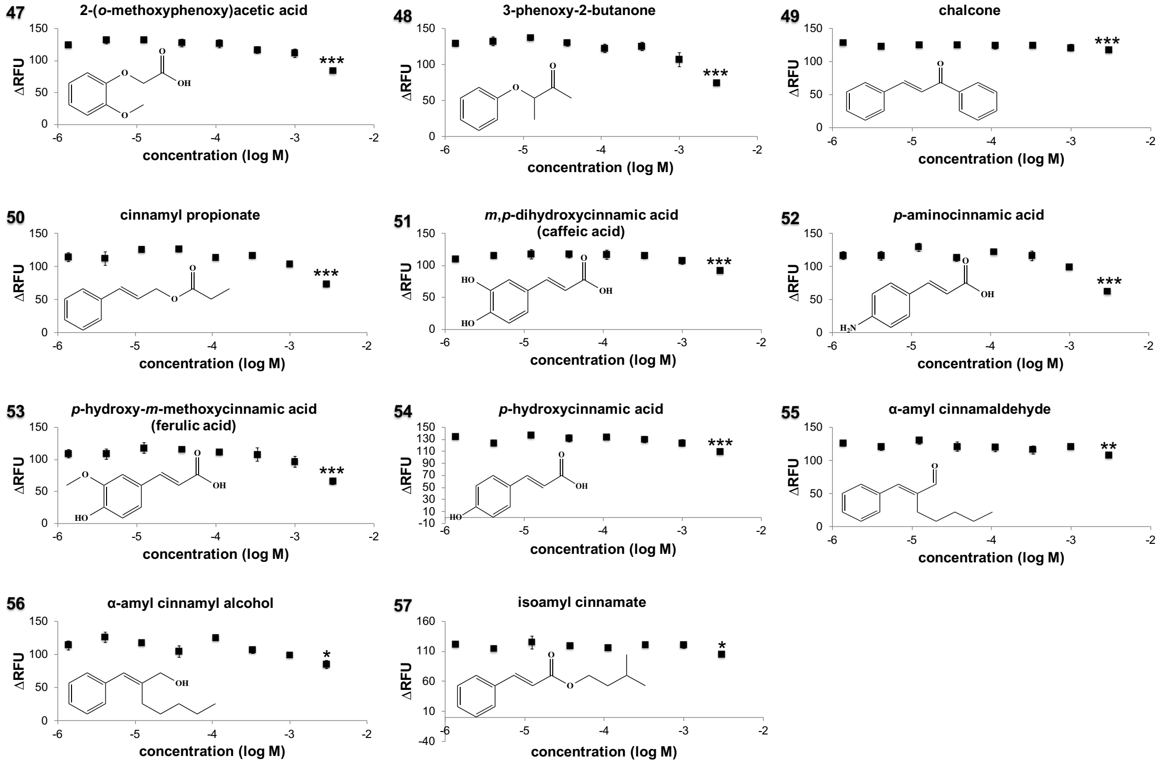


#### Neutral

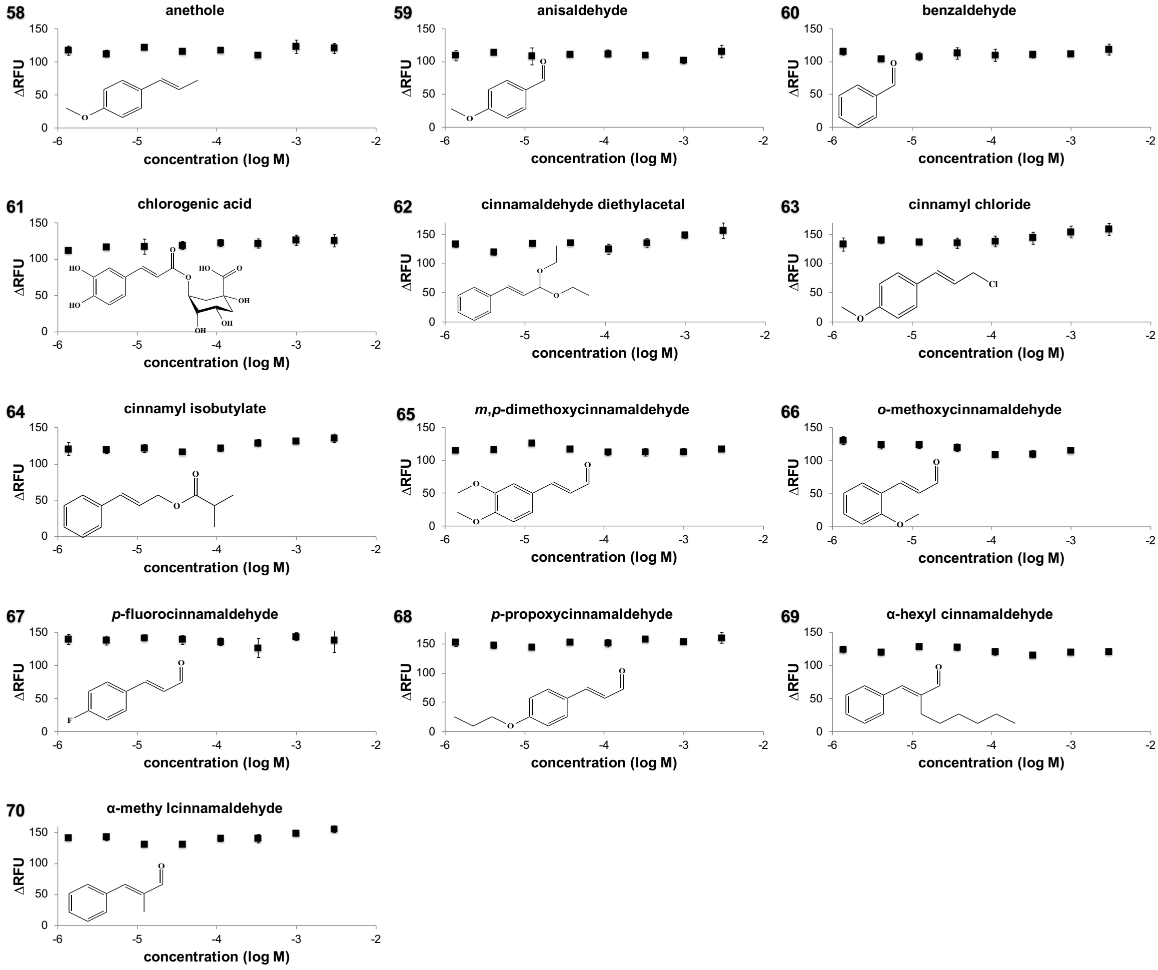

### supplemental Figure 6

#### Positive allosteric threshold -2.5

**71** ***p*-hydroxycinnamaldehyde**

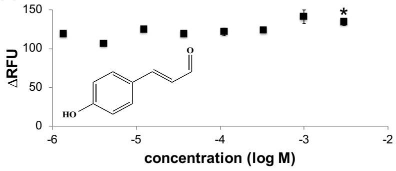


#### Positive allosteric threshold -3.0

**72** **p-ethoxybenzaldehyde**

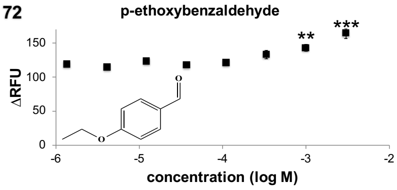


**73** **phenylpropargyl aldehyde**

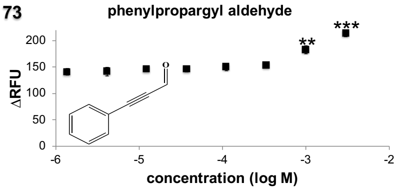


**74** **cinnamaldehyde oxime**

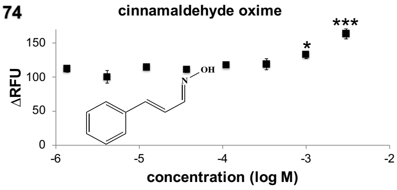


#### Positive allosteric threshold -4.0

**75** **cinnamonitrile**

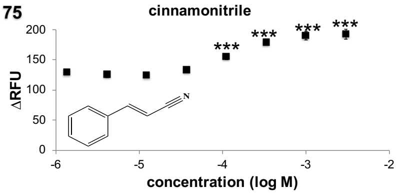


**76** **cinnamaldehyde (CA)**

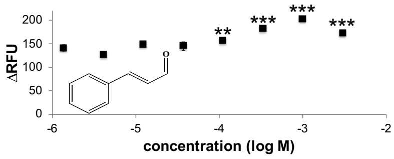


**77** ***p*-ethoxycinnamaldehyde**

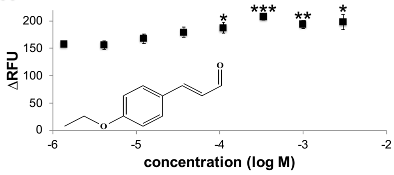


**78** ***p*-methoxycinnamonitrile**

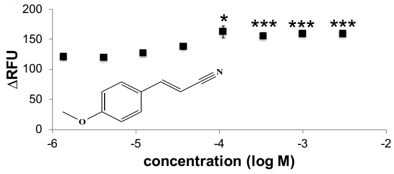


#### Positive allosteric threshold -4.4

**79** **p-methoxycinnamaldehyde**

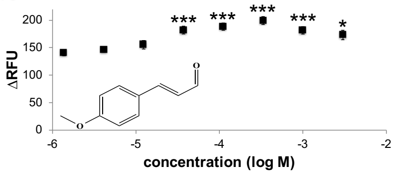

supplemental Figure 7

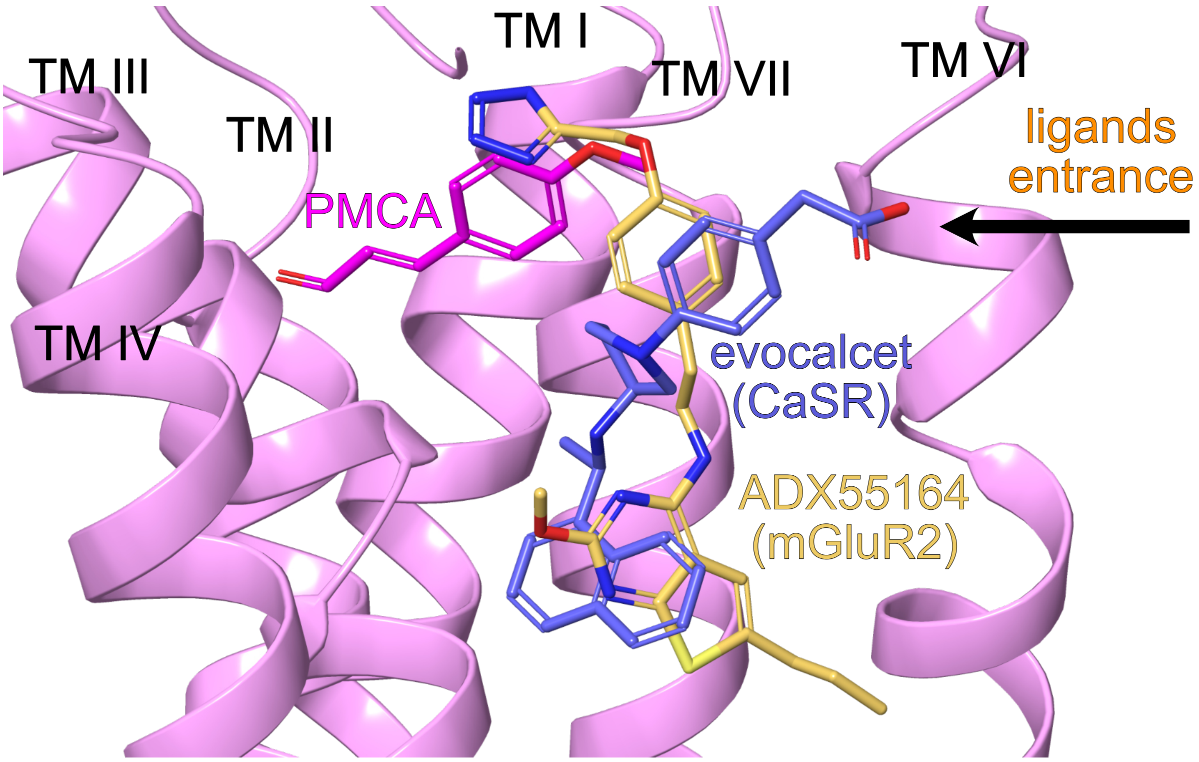
